# Targeting of SUMOylation leads to cBAF complex stabilization and disruption of the SS18::SSX transcriptome in Synovial Sarcoma

**DOI:** 10.1101/2024.04.25.591023

**Authors:** Konstantinos V. Floros, Carter K. Fairchild, Jinxiu Li, Kun Zhang, Jane L. Roberts, Richard Kurupi, Bin Hu, Vita Kraskauskiene, Nayyerehsadat Hosseini, Shanwei Shen, Melissa M. Inge, Kylie Smith-Fry, Li Li, Afroditi Sotiriou, Krista M. Dalton, Asha Jose, Elsamani I. Abdelfadiel, Yanli Xing, Ronald D. Hill, Jamie M. Slaughter, Mayuri Shende, Madelyn R Lorenz, Mandy R. Hinojosa, Benjamin R. Belvin, Zhao Lai, Sosipatros A. Boikos, Angeliki M. Stamatouli, Janina P. Lewis, Masoud H. Manjili, Kristoffer Valerie, Renfeng Li, Ana Banito, Andrew Poklepovic, Jennifer E. Koblinski, Trevor Siggers, Mikhail G. Dozmorov, Kevin B. Jones, Senthil K. Radhakrishnan, Anthony C. Faber

## Abstract

Synovial Sarcoma (SS) is driven by the SS18::SSX fusion oncoprotein. and is ultimately refractory to therapeutic approaches. SS18::SSX alters ATP-dependent chromatin remodeling BAF (mammalian SWI/SNF) complexes, leading to the degradation of canonical (cBAF) complex and amplified presence of an SS18::SSX-containing non-canonical BAF (ncBAF or GBAF) that drives an SS-specific transcription program and tumorigenesis. We demonstrate that SS18::SSX activates the SUMOylation program and SSs are sensitive to the small molecule SAE1/2 inhibitor, TAK-981. Mechanistically, TAK-981 de-SUMOylates the cBAF subunit SMARCE1, stabilizing and restoring cBAF on chromatin, shifting away from SS18::SSX-ncBAF-driven transcription, associated with DNA damage and cell death and resulting in tumor inhibition across both human and mouse SS tumor models. TAK-981 synergized with cytotoxic chemotherapy through increased DNA damage, leading to tumor regression. Targeting the SUMOylation pathway in SS restores cBAF complexes and blocks the SS18::SSX-ncBAF transcriptome, identifying a therapeutic vulnerability in SS, positioning the in-clinic TAK-981 to treat SS.

## Introduction

Synovial sarcoma (SS) is an aggressive soft tissue sarcoma (STS) that occurs frequently in pediatric and young adults. Metastatic SS remains incurable with little benefit observed even with newer combination chemotherapies (Burningham et al., 2012). Given the lack of durable responses to chemotherapies, development of other strategies and targeted therapies for SS has been a priority. Coding for 78 amino acids from the C-terminal of SSX1 or SSX2 or SSX4, each found on the X chromosome is fused in frame for coding with amino acids 1–379 of the SS18 subunit, generating SS18::SSX1, or SS18::SSX2, or SS18::SSX4, each of which is pathognomonic for SS (Clark et al., 1994). Genetic inhibition of SS18::SSX results in growth arrest and cell death in SS cells *in vitro* (Carmody Soni et al., 2014) and inhibition of tumor growth *in vivo* (Takenaka et al., 2010). The SS18::SSX fusion oncoprotein is dominant and there appears to be infrequent secondary genetic events in SS(McBride et al., 2018). The known reliance of SS on SS18::SSX to drive and maintain tumorigenicity makes SS18::SSX an appealing therapeutic target; however, direct small-molecule inhibitors or degraders for SS18::SSX have yet to be developed(McBride *et al*., 2018).

SS18::SSX includes no typical DNA-binding doman (Tamaki et al., 2015), but does act as a transcriptional modifier (Zöllner et al., 2015). WT SSX is thought to function in the repression of gene transcription, likely through polycomb repressive complex interactions 1 and 2, but its function has not been deeply explored (Boulay et al., 2021). WT SS18 is a dedicated member of the chromatin remodeling mammalian SWI/SNF (mSWI/SNF, also called BAF) complex (McBride *et al*., 2018; Zöllner *et al*., 2015). In humans, there are several BAF complex subtypes, each consisting of 13 or more subunits of both shared and unique members (Kadoch et al., 2013). The major BAF complexes in most cells include canonical BAF (cBAF, also referred to as BAF), (polybromo-associated BAF), PBAF, and non-canonical or GLTSCR1-containing (ncBAF or GBAF). Each of these three complexes impacts gene transcription uniquely and, depending on the context, can both positively and negatively regulate gene expression. In recent years, it has become apparent that the oncogenicity of SS18::SSX can largely be attributed to the aberrant interaction of SS18::SSX with BAF complexes (Brien et al., 2018; Li et al., 2021a; McBride *et al*., 2018; Michel et al., 2018).

Like WT SS18, SS18::SSX can stably incorporate into BAF complexes; when this happens, it competes with and leads to the degradation of SS18 (Kadoch and Crabtree, 2013), In addition, we have recently demonstrated that SS18::SSX leads to whole complex cBAF degradation, the result of which is increased relative prevalence of other BAF-family subtypes, in particular ncBAF with the fusion, altering the cellular composition from a dominant cBAF phenotype to a mix of ncBAF, PBAF and cBAF(Li *et al*., 2021a). Furthermore, overexpression of SMARCB1 is sufficient to restore cBAF genome-wide occupancy and to block the growth of SS cells (Nakayama et al., 2017). Disruption of the ncBAF-only components, BRD9 or GLTSCR1 abrogates ncBAF function and SS cell growth (Brien *et al*., 2018; Michel *et al*., 2018). Thus, SS is defined by a rebalancing of BAF complexes on chromatin leading to an ncBAF dominant transcriptome, the reversal of which by restoration of cBAF complexes and/or depletion of ncBAF complexes is toxic to SS.

SUMOylation is a highly conserved eukaryotic post-translational modification (PTM) of proteins that results in the covalent attachment of SUMO (<u>s</u>mall <u>u</u>biquitin-like <u>mo</u>difier) to lysine residues of the target protein. SUMOylation functions in a wide array of biological processes. These include chromatin modeling, transcription, repair of DNA, cell death, and metabolism (Celen and Sahin, 2020; Hay, 2005; Li et al., 2021b). Of note, there is very little known about SUMOylation in SS (Sun et al., 2011).

Subasumstat (TAK-981) is a first-in-class SUMOylation inhibitor that has recently been demonstrated to disrupt SAE (SUMO-Activating Enzyme) function by covalently binding to SUMO resulting in the formation of an adduct (Langston et al., 2021). TAK-981 has advanced through phase I trials in patients with late-stage cancers where a recommended phase 2 dose (RP2D) was established (NCT03648372). TAK-981 is also in a phase I trial in combination with pembrolizumab (NCT04381650) in metastatic solid tumors. Thus, TAK-981 allows for clinical evaluation of SUMOylation inhibition as a tractable pathway in cancer.

## Results

### Synovial sarcoma is highly sensitive to disruption of the SUMOylation pathway

To explore the reliance of SS on SUMOylation for survival, we interrogated the Broad Institute depository (DepMap) of genome-wide RNAi screens across more than 800 cancer cell lines. Strikingly, SS was the most sensitive subtype of cancer to knocking down *UBE2I* (*UBC9*), *SUMO2* (the most widely expressed SUMO in humans (Salas-Lloret and Gonzalez-Prieto, 2022)), *PIAS1*, *SAE1 and UBA2* (*SAE2*) (Fig. S1A-E), molecules encompassing the entire SUMOylation process (Fig. S1F) and pointing to a selective dependency of SS on this PTM. Indeed, analysis of the effective RNAi in the screen against a 12 gene ‘’SUMO signature’’ created from the Biocarta SUMO pathway (Hoellein et al., 2014), corroborated pathway sensitivity (Fig. 1A). Next, we determined whether the expression of SUMOylation pathways proteins in SS depends on the presence of SS18::SSX. For this, we parsed data from a recent study (McBride *et al*., 2018) and e evaluated the mRNA presence of key members of the SUMOylation cycle (Fig. S1F), following knockdown of short hairpins against the fusion oncogene (shSSX). Essential components of the SUMOylation pathway were significantly reduced after silencing SS18::SSX (Fig. 1B), demonstrating a role of SS18::SSX to increase SUMOylation in SS.

**Figure 1.**
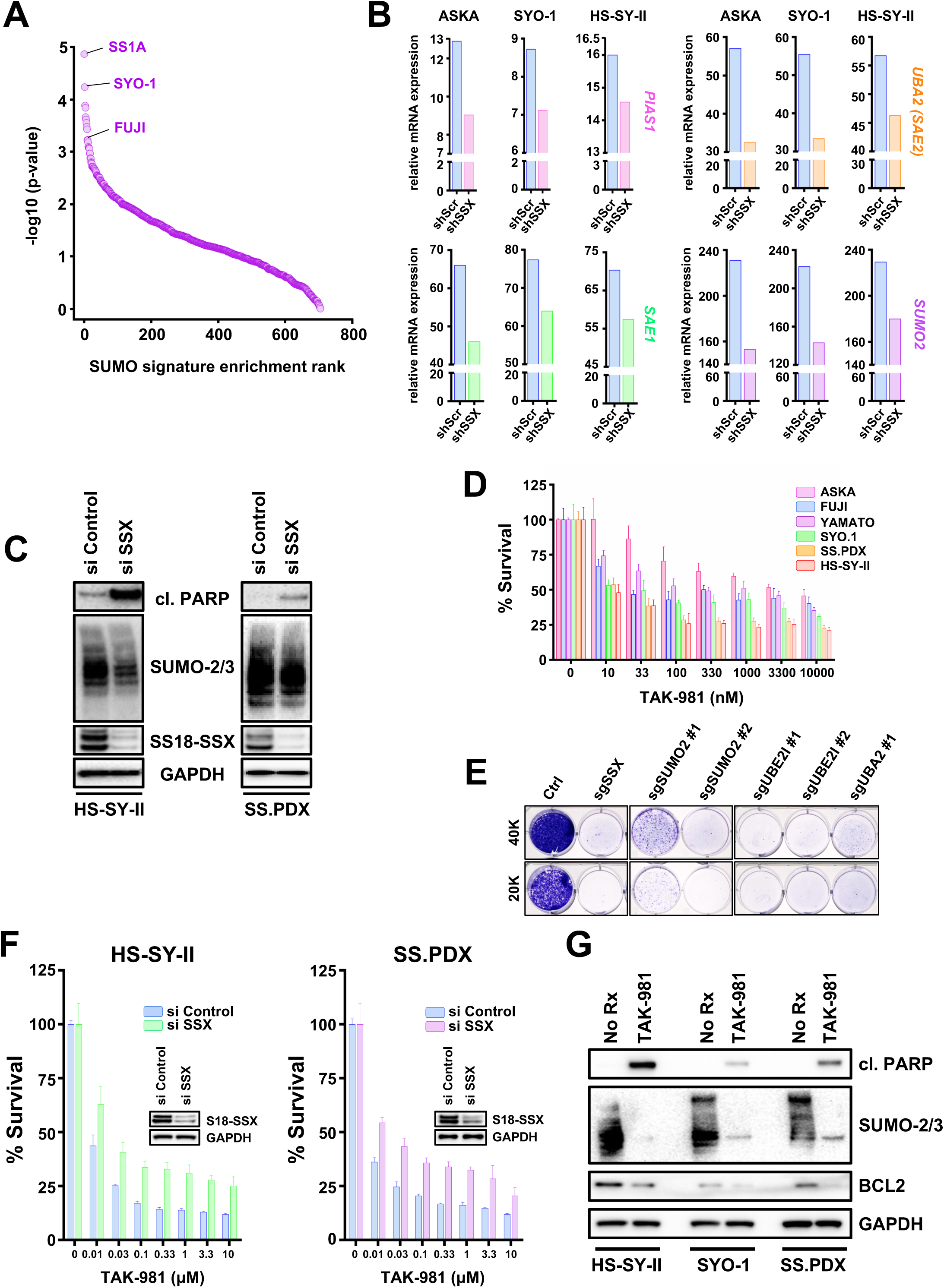
**A)** Cell lines ranked by enrichment (-log10(p-value)) in the SUMO signature (BIOCARTA_SUMO_PATHWAY) using single-sample GSEA on cell line-specific genes ordered by the gene dependency scores (DEMETER2 scores from DepMap). **B)** RNA levels of genes involved in SUMOylation following shSSX knockdown were obtained from a RNA seq analysis performed in SS cell lines **C)** SS cells were transfected with si SSX for 18h and whole cell lysates (supplemented with 0.5M of N-ethylmaleimide (NEM)) were probed with the indicated antibodies. **D)** SS cells were treated with increasing concentrations of the SUMOylation inhibitor, TAK-981, and cell viability was assessed 72h later using cell-titer glo. **E)** HS-SY-II cells underwent CRISPR/Cas9-mediated gene targeting of the indicated SUMOylation genes and cells were counted and plated at low density (20.000 or 40.000 cell per 6 -well plate) and stained with crystal violet 12 days later. **F)** SS cells were transfected with siRNA directed against SSX for 36h, reseeded, and treated with increasing concentrations of TAK-981 for 72h before cell viability was assessed (inset is immunoblotting confirming knockdown). **G)** SS cells were treated with 100nM TAK-981 for 36h or left untreated (No Rx) and whole cell lysates (supplemented with 0.5M of N-ethylmaleimide (NEM)) were probed with the indicated antibodies.

We corroborated these data by silencing SS18::SSX in HS-SY-II cells (Sonobe et al., 1992) and the *ex vivo* cell line, SS.PDX, derived from a SS-patient-derived xenograft (PDX) model (Fairchild et al., 2021). SUMO modified proteins decreased in both cell lines, which also underwent cell death (as evidenced by cleaved PARP) following depletion of SS18::SSX (Fig. 1C). Additionally, markedly reduced SUMOylation was demonstrated in a third SS cell line, Yamato (Naka et al., 2010), subjected to inducible knock down of SS18::SSX (Fig. S2A). Altogether, these data demonstrate SS18::SSX increases SUMOylation in SS.

To verify the pathway dependency on SUMOylation in SS, we pharmacologically blocked global SUMO modifications, utilizing the novel SUMOylation inhibitor, TAK-981 (subasumstat) (Langston *et al*., 2021). The cell viability of the SS cell lines tested was robustly reduced after treatment with increasing concentrations of TAK-981, as demonstrated by CellTiter-Glo and crystal violet survival assays (Fig. 1D and S2B). To further corroborate the dependence of SS on SUMOylation, we depleted characteristic components of the SUMO pathway using sgRNAs in the HS-SY-II cells. Genetically targeting of SUMOylation resulted in nearly complete death of the SS cells, as evidenced by crystal violet (Fig. 1E). To determine whether the toxicity of TAK-981 in SS was at least partially due to SS18::SSX, we transfected two SS cell lines, HS-SY-II and SS.PDX *ex vivo* cells, with siRNA targeting SS18::SSX, then treated them with increasing concentrations of TAK-981 and performed CellTiter-Glo viability assays. A significant rescue of TAK-981 induced toxicity by SS18::SSX depletion was noted in both cell lines (Fig. 1F). Enhanced cleavage of PARP was detected in TAK-981 treated HS-SY-II, SS.PDX and SYO-1 cells (Langston *et al*., 2021), demonstrating a cell death effect. Of note, downregulation of the well-established histological marker of SS, the prosurvival BCL2 protein (Barrott et al., 2017; Fairchild *et al*., 2021; Hirakawa et al., 1996), was also observed following TAK-981 treatment, without changes in other BCL2 family member proteins (Fig. 1G and S2C). Interestingly, we have shown BCL2 is upregulated in the presence of SS18::SSX in human and mouse SS models (Jones et al., 2013), suggesting to us that TAK-981 may impact the SS18::SSX-driven transcriptome.

### TAK-981 de-SUMOylates SMARCE1, an integral subunit of the cBAF and PBAF complex

We next sought to gain mechanistic insights into why blocking SUMOylation in SS is so effective. To identify the members of SS proteome that were de-SUMOylated upon TAK-981 treatment, a proteome-wide detection of the SUMOylation sites (Lumpkin et al., 2017) in HS-SY-II SS cells before and after TAK-981 treatment (Fig. 2A) was performed. The assay utilizes a wild-type α-lytic protease (WaLP) to digest SUMOylated proteins resulting in the production of peptides carrying SUMO-remnant diglycyl-lysine (KGG) at the site of SUMO modification (Cell Signaling technology). Using specific antibodies for the isolation of the KGG-containing peptides and followed by mass spectrometry for sequencing of the captured peptides, we identified 118 SUMO-modified proteins in the control HS-SY-II cells that were significantly reduced by TAK-981 (Tab. S1). Among these, was the cBAF complex member, SMARCE1. SMARCE1 constitutes a fundamental component of the BAF core module along with SMARCC1, SMARCC2, SMARCD1/2/3 and SMARCB1(Mashtalir et al., 2018). SMARCE1 incorporates into cBAF and PBAF, but not ncBAF (Mashtalir *et al*., 2018). Importantly, it has been recently demonstrated that in clear cell meningioma (CCM), driven by loss of SMARCE1, the cBAF complex fails to stabilize on chromatin, attenuating its activity (St Pierre et al., 2022).

**Figure 2.**
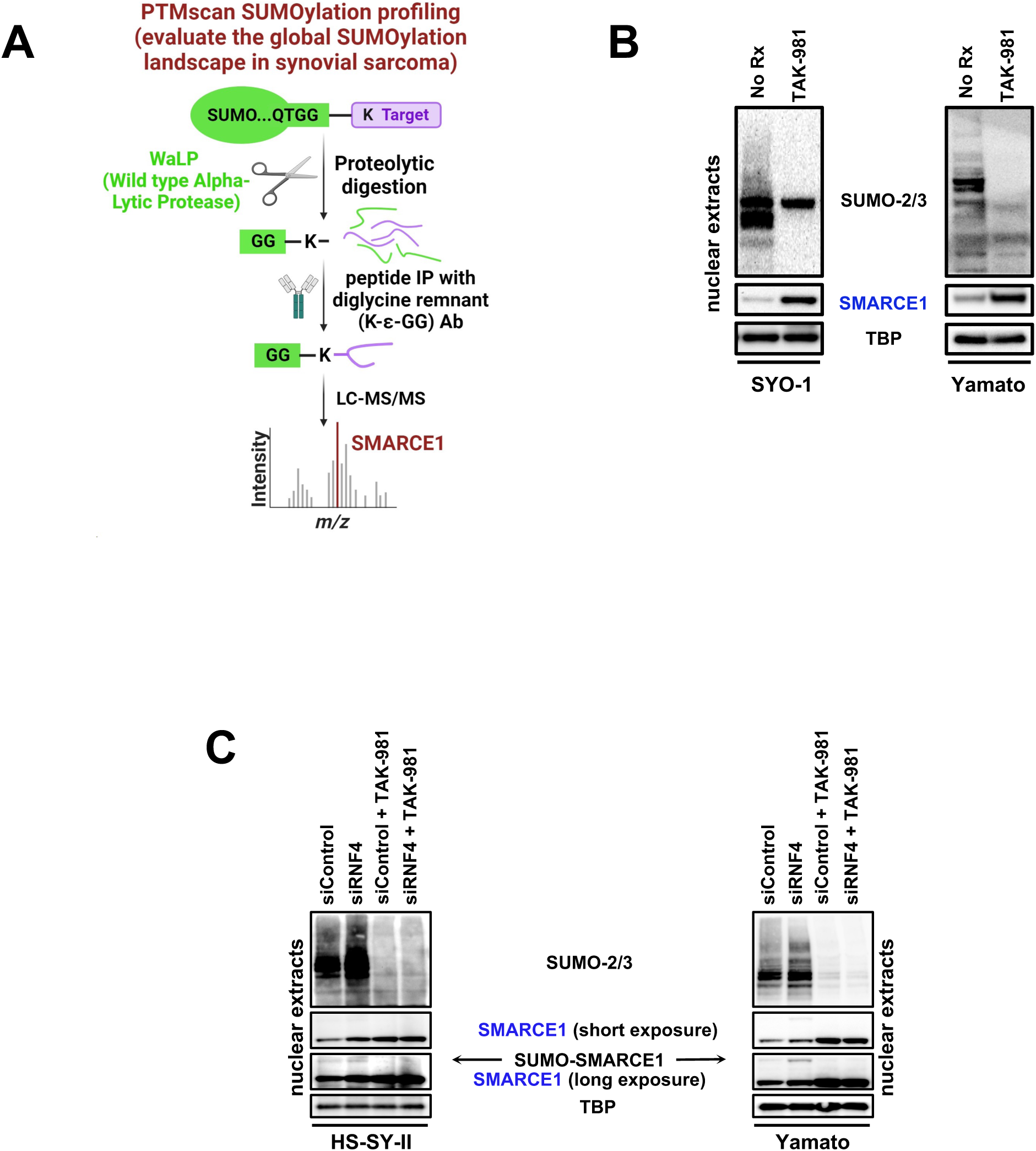
**A)** Illustration depicting the method followed for the Proteome-wide identification of SUMO modification sites by mass spectrometry (PTMscan SUMOylation profiling) of the HS-SY-II cells treated either with a) No Rx or b) 100 nM of TAK-981 for 36h prior to analysis. **B)** SS cells were treated with 100nM TAK-981 for 36h or left untreated (No Rx) and nuclear lysates (supplemented with 0.5M of N-ethylmaleimide (NEM)) were probed with the indicated antibodies. **C)** and **D)** SS cells transfected with scramble (sc) siRNA or siRNA directed against siRNF4 were treated with 100nM TAK-981 for 36h or left untreated (No Rx) and nuclear lysates (supplemented with 0.5M of N-ethylmaleimide (NEM)) were probed with the indicated antibodies. TBP was used as a nuclear loading control.

We therefore determined the effect of TAK-981 on SMARCE1 protein levels. Interrogation of SYO-1 and Yamato cells by immunoblotting after TAK-981 addition displayed a striking upregulation of SMARCE1 following de-SUMOylation (Fig. 2B). Consistent with the literature that the attachment of SUMO moieties can prime the target protein for ubiquitylation by the E3 ubiquitin ligase, RNF4 (Sun et al., 2007), and subsequent proteasome degradation, we found that knocking down RNF4 by siRNA leads to a clear upregulation of the SMARCE1 protein in the easier transfected SS cell lines, HS-SY-II and Yamato cells (Fig. 2C). Importantly, RNF4 depletion resulted in accumulation of the SUMO-conjugated SMARCE1 protein no longer degraded by the proteasome, as indicated by the appearance of a band with a size consistent with the SUMOylated form of the protein, that was undetectable in the presence of TAK-981 (Fig. 2C). As expected, SMARCE1 protein accumulated following TAK-981 treatment (Fig. 2C), but no more so upon depletion of RNF4 in the presence of TAK-981. Altogether, these data demonstrate that SUMOylation inhibition blocks RNF4-mediated degradation of the SMARCE1 cBAF subunit.

### SUMOylation inhibition stabilizes the cBAF complex

The role of SMARCE1 in sustaining the integrity of the cBAF complex, by bolstering the connectivity between the unique cBAF component, ARID1A, and core module constituents, like SMARCB1, has already been described(St Pierre *et al*., 2022). Supporting the hypothesis of cBAF complex stabilization following TAK-981 treatment and subsequent SMARCE1 upregulation, we observed an increase in fellow cBAF members ARID1A, and SMARCB1 (BAF47) in SYO-1 and HS-SY-II cells in the presence of TAK-981 (Fig. 3A). Seeking to understand the consequence of upregulated SMARCE1 in relation to the three different BAF complexes in SS, we determined the alterations in the size of the BAF complexes by conducting a density sedimentation study. Interrogation of glycerol gradients of the nuclear extracts revealed a marked upshift of cBAF members SMARCE1, ARID1A and BAF47 by approximately 3 fractions after TAK-981 treatment in SYO-1 and HS-SY-II cells (Fig. 3B and 3C), in line with the expected gain in complex mass resulting from the incorporation of larger amounts of SMARCE1, ARID1A and BAF47. There were also modest increases in pBAF stability (Fig. 3B and 3C). There were no (SYO-1 cells) or slight (HS-SY-II cells) increases in ncBAF stability (Fig. 3B and 3C). Overall, these data demonstrate that SMARCE1 protection from SUMOylation-mediated degradation stabilizes cBAF complexes and to a lesser extent, pBAF complexes.

**Figure 3.**
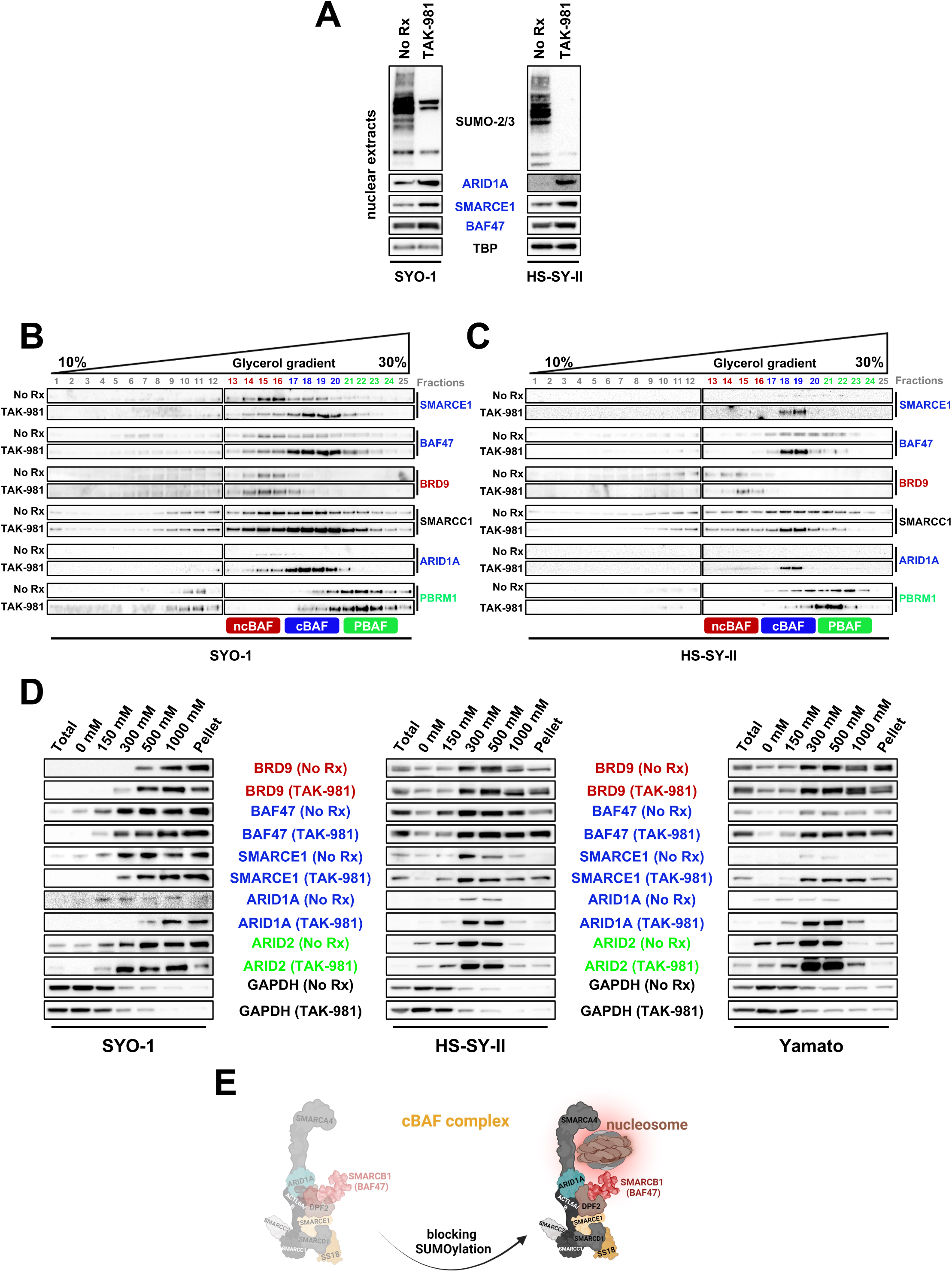
**A)** SS cells were treated with 100nM TAK-981 for 36h or left untreated (No Rx) and nuclear lysates (supplemented with 0.5M of N-ethylmaleimide (NEM)) were probed with the indicated antibodies. TBP was used as a nuclear loading control. **B) and C)** SYO-1 and HS-SY-II cells were pretreated with no drug or 100 nM of TAK-981 for 36h. Density sedimentation assay (10%–30% glycerol gradient) followed by immunoblotting the indicated BAF family components was performed on nuclear extracts. **D)** Immunoblot of SS cells for specific BAF components was performed following treatment with no drug and 100 nM of TAK-981 for 36h and differential salt extraction (0–1,000 mM NaCl). GAPDH was used as a cytoplasmic loading control. **E)** Illustration depicting the changes in chromatin affinity for the cBAF complex after TAK-981 treatment.

While these data demonstrated enhanced protein complexing, the assays did not speak to any changes of these complexes at the chromatin. We therefore measured relative chromatin affinities before and after TAK-981 treatment, performing differential salt extraction in both SS cell lines. The higher affinity of the cBAF complex for chromatin after rescue of SMARCB1/BAF47 has been delineated in the past (Michel *et al*., 2018). Blocking SUMOylation resulted in cBAF complex dissociation from chromatin at higher salt concentrations (300-500 mM NaCl) than prior to drug treatment, as evidenced by ARID1A, BAF47 and SMARCE1 expression levels (Fig. 3D). In contrast, the elution of BRD9 was noted at lower NaCl concentrations following TAK-981 treatment in SYO-1 and Yamato cells. These results suggest a stronger binding of the cBAF complex to chromatin after SUMOylation inhibition and a relative shift away from the dominance of SS18::SSX-ncBAF complexes at the chromatin in SS (Fig. 3E).

### Blocking SUMOylation leads to disruption of the synovial sarcoma signature and to induction of mesenchymal differentiation

Results of a previous study by Banito et al.(Banito et al., 2018) demonstrated that SS18::SSX binds to and activates the expression of KDM2B-PRC1 target genes thereby supporting synovial sarcomagenesis and ultimately defining the SS signature. This signature primarily contains genes that encode homeobox transcription factors related to neurogenesis and other developmental processes (Fig. S4B). SS18::SSX knockdown reduces this SS signature and rewires the cells towards a mesenchymal phenotype, including the expression of genes encoding extracellular matrix (ECM) proteins and secreted proteins that are highly expressed in human fibroblasts. We evaluated whether the addition of TAK-981 can efficiently impede the activation of the SS18::SSX transcriptome and hence disrupt the SS signature, while inducing ECM-related genes, in total resembling the changes observed after silencing the fusion protein. Due to the heterogeneity of the gene targets of SS18::SSX in SS, we performed RNA-seq in three SS cell lines (SYO-1, the *ex vivo* SS.PDX, and HS-SY-II), before and after treatment with TAK-981. Strikingly, the up- and down-regulated pathways, based on the expression changes of the related genes, verified our hypothesis as several of the most enhanced or suppressed biological processes are common to both TAK-981-treated SS cells and SS18::SSX-genetically inhibited SS cells (Fig. 4A). In addition, there is a significant overlap of the altered genes between TAK-981-treated cell lines (Fig. S3A and S3B), which was verified also after pairwise comparisons (Fig. S3C, S3D and S3E). Consistent with the pathway analyses (Fig. 4A), the overlap between the genes up- and down-regulated after knocking down SS18::SSX in HS-SY-II cells (Banito *et al*., 2018), and the genes enhanced or suppressed after the addition of TAK-981 to SYO-1 and SS.PDX cells were significant (Fig. 4B and 4C). Noteworthy, this correlation was even more significant when comparing the SS18::SSX-genetically inhibited HS-SY-II cells and TAK-981-treated HS-SY-II cells (Fig. 4B). Consistently, a significant portion of the 1000 most up- or down-regulated genes after TAK-981 treatment are also up- or down-regulated after silencing SS18::SSX in all three SS cell lines (Fig. 4D). Distribution of the commonly altered genes according to their fold change revealed a positive correlation when interrogating the cell lines in pairwise fashion (Fig. S3F, S3G, S3H), verifying their uniform response to the inhibition of SUMOylation in SS. Additionally, addition of TAK-981 promotes silencing of developmental genes and augments the expression of genes related to ECM, cell adhesion and muscle function (Fig. S3I and S3K), similar to the changes observed after SS18::SSX silencing, and in line with the gene ontology conducted in Fig. 4A. The above data were further corroborated by qPCR (Fig. S3J and S3L) for most of the gene changes identified by RNA-seq.

**Figure 4.**
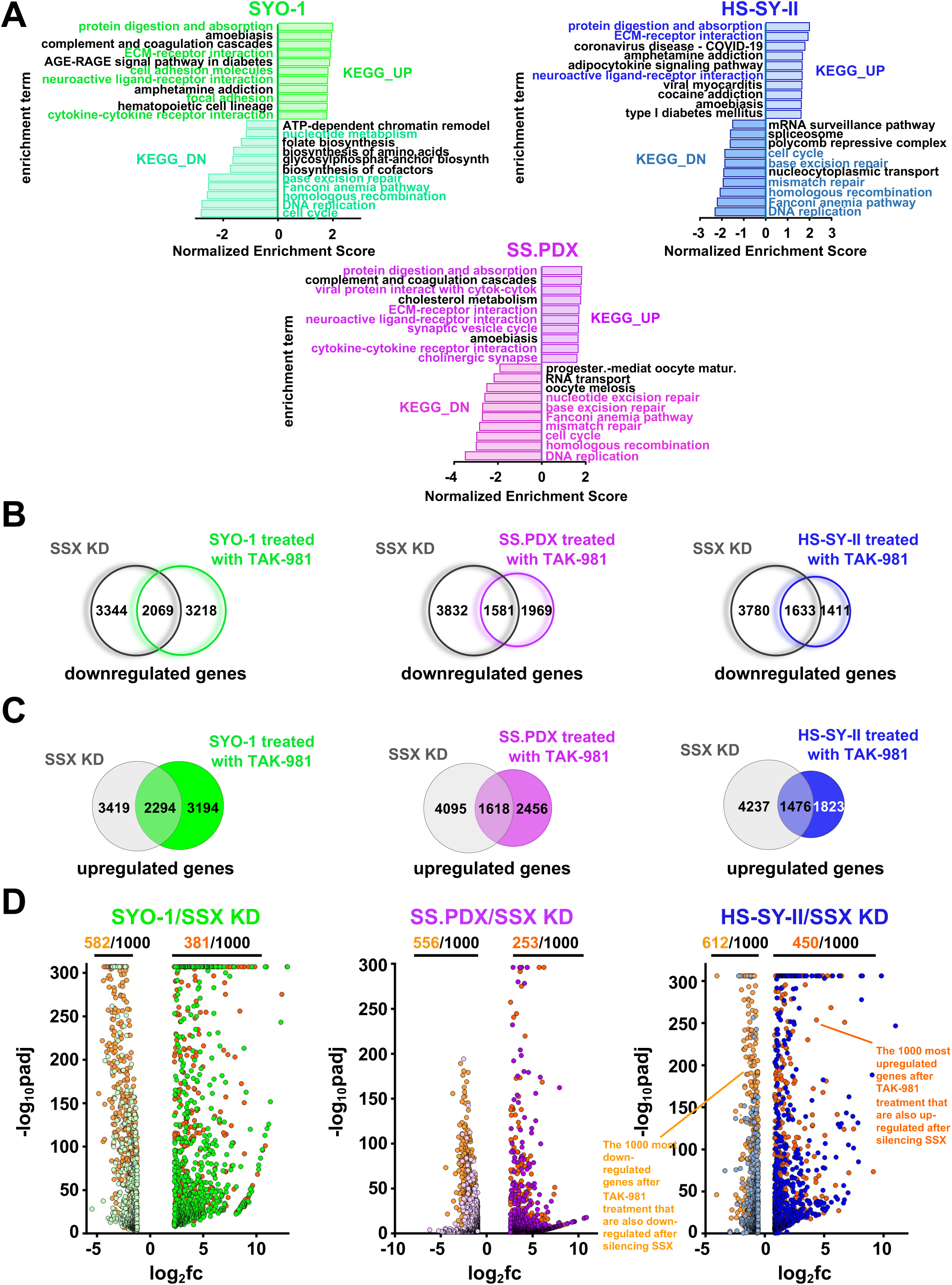
**A)** Normalized enrichment scores after KEGG analysis of RNA-seq data demonstrating pathways altered by 100nM TAK-981 treatment for 36h in the indicated SS cells. **B)** Venn diagram of genes significantly downregulated following SS18-SSX knockdown or treatment with 100nM TAK-981 for 36h. **C)** Venn diagram of genes significantly upregulated following SS18-SSX knockdown or treatment with 100nM TAK-981 for 36h. **D)** Volcano plots of the most down- or upregulated 1000 genes following SS18-SSX knockdown plotted with the common downregulated genes (orange dots) or common upregulated genes (red dots) by RNA seq, among the indicated SS cell line.

### The binding of BAF complexes to chromatin is responsible for many of the observed expression changes of the SS transcriptome after blocking SUMOylation

To further investigate how the different BAF complexes contribute to the observed transcription phenotype following suppression of SUMOylation, we performed Chromatin Immunoprecipitation (ChIP)-seq using antibodies raised against SMARCA4 (BRG1), the ATPase found in all three BAF complexes, and against the fusion protein SS18::SSX (a member of ncBAF). Gene ontology analysis of the reduced and the enhanced SMARCA4 ChIP seq signals after TAK-981 treatment in HS-SY-II cells indicated transcriptome changes of genes intimately involved in neurogenesis, DNA replication and cell cycle (for the reduced signals) and genes involved in the ECM and muscle function (for the enhanced signals) (Fig. 5A and 5D), indicating a correlation between the RNA-seq data and the binding profiles of all three BAF complexes. Moreover, more than half of the genes with a decreased SMARCA4 ChIP-seq signal (155 out of 288) also displayed a reduced SS18-SSX ChIP-seq signal (Fig. 5B, 5F). Additionally, 49 out of the 155 genes that had enhanced SS18-SSX signals also had enhanced SMARCA4 binding (Fig. 5E, 5F, Fig. S4C and S4E).Furthermore, 15 genes that demonstrated significant downregulation of SS18::SSX binding (Fig. 5B) (Banito *et al*., 2018), and 24 genes with significant downregulation of SMARCA4 binding, were included among the 100 most downregulated genes following SS18::SSX knock down (Fig. 5B). While we did not see significant overlap between increased SMARCA4 or SS18::SSX binding at the genes that were upregulated after silencing SS18::SSX (Banito *et al*., 2018), binding of SMARCA4 at several genes related to the mesenchymal phenotype that were upregulated following SS18::SSX silencing was noted (Fig. 5G). Next, when we examined the overlap between the up- and down-regulated genes following TAK-981 treatment in HS-SY-II cells - identified by the RNA seq analysis - and the SMARCA4/SS18::SSX binding profiles, we again found significant overlap (Fig. S4A and S4D), consistent with the comparisons to the gene expression changes following SS18::SSX knock down. Additionally, a significant portion of the genes that demonstrated reduced KDM2B binding demonstrated reduced SMARCA4/SS18::SSX binding after TAK-981 treatment (Fig. S4A). We then turned our attention to a 30-gene signature that was developed by examining genes specifically elevated in SS compared to other sarcoma tumors and that were diminished after silencing the fusion protein in HS-SY-II cells; these genes constitute a core ‘’SS signature’’(Banito *et al*., 2018). More than half of these genes had weaker SMARCA4 and/or SS18::SSX binding after TAK-981 treatment (Fig. S4A and S4B).

**Figure 5.**
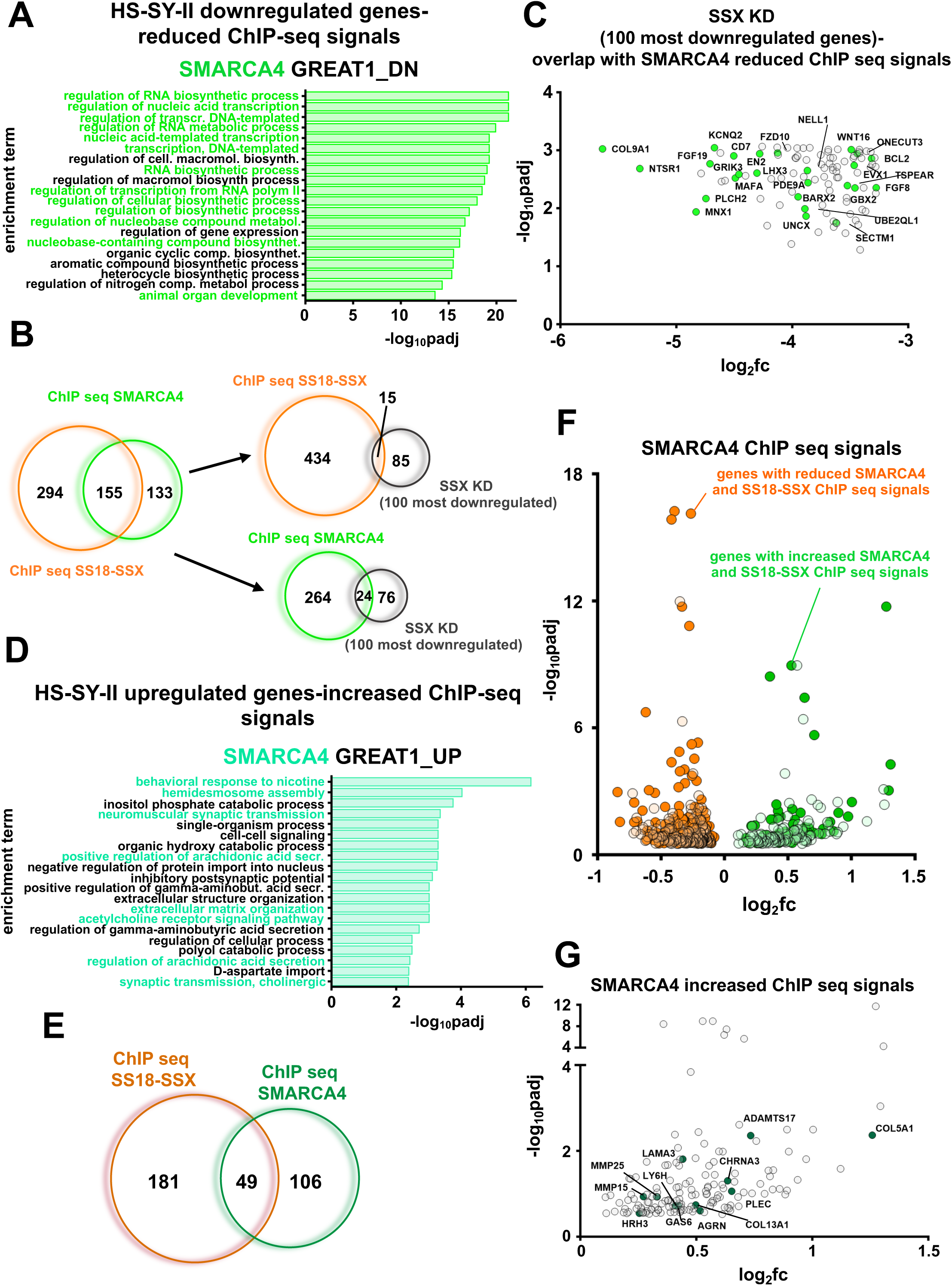
**A)** Gene ontology analysis of genes associated with significantly reduced SMARCA4 ChIP-seq signals after TAK-981 treatment in HS-SY-II cells. **B)** Venn diagrams of the overlap in significantly downregulated genes among the different noted experiments. **C)** Overlap of 100 most significantly downregulated genes following SS18-SSX knockdown with significantly reduced SMARCA4 ChIP seq signals. **D)** Gene ontology analysis of genes associated with significantly upregulated SMARCA4 ChIP-seq signals after TAK-981 treatment in HS-SY-II cells. **E)** Venn diagram of the overlap in significantly upregulated genes among the different noted experiments. **F)** Overlap of the genes with both reduced SMARCA4 ChIP-seq signals and SS18-SSX ChIP-seq signals (orange dots), and genes with both increased SMARCA4 ChIP-seq signals and SS18-SSX ChIP-seq signals (green dots). **G)** Genes related to the mesenchymal phenotype associated with increased SMARC4A ChIP-seq signals (green dots).

We next evaluated genes with significantly enhanced or diminished SMARCA4 and SS18::SSX ChIP-seq peaks following TAK-981 treatment with that of the H3K27ac ChIP-seq, a marker for active gene enhancers and promoters (Shlyueva et al., 2014). Noteworthy, the attenuated ChIP-seq profile of SS18::SSX and of H3K27ac at genes overlaid significantly with each other (Fig. S4A), as did the enhanced ChIP-seq profile of SMARCA4 and of H3K27ac (Fig. S4D and S4F). We next analyzed ChIP-seq profiles of SS18-SSX, KDM2B, and H3K27ac in SYO-1 cells, before and after TAK-981 treatment. Similar to the data in HS-SY-II cells, we found significant overlap in the changes in ChIP-seq signals with TAK-981 treatment among these different experiments, as well as with these changes and the changes in gene expression following SS18-SSX knockdown in the HS-SY-II cells (Banito *et al*., 2018) or TAK-981 treatment, as well as the SS signature (Banito *et al*., 2018) (Fig. S5A-S5K). Altogether, the above data suggests that while reduced ncBAF complex plays a central role in the loss of the SS18::SSX-driven transcriptome, the changes noted in other BAF complexes (for instance, increased cBAF binding and activity) contributes to the transcriptome changes following SUMOylation inhibition in SS.

### Addition of TAK-981 results in loss of chromatin accessibility to promoters of the SS18::SSX transcriptome

SS18::SSX inhibition in SS causes a large reduction of chromatin accessibility at SS18::SSX binding sites, associated with rearrangement of BAF complexes across the genome(Banito *et al*., 2018) Based on the data above, we reasoned that inhibition of SUMOylation resulted in mitigated accessibility to the promoters of genes in the the SS18::SSX transcriptome. Therefore, we performed ATAC-seq in HS-SY-II and SYO-1 cells before and after TAK-981 treatment. Subsequently, we focused on the reduced ATAC-seq signals at the genes that were also suppressed in HS-SY-II cells after knocking down SS18::SSX (Banito *et al*., 2018). Strikingly, blocking SUMOylation caused a marked reduction of the accessibility in SYO-1 cells, and a smaller but clear attenuation of the signal in the HS-SY-II cells, at the promoters of the genes reduced in expression following SS18::SSX knockdown. In addition, we observed enhancement at introns and at distal intergenic sites of these genes (Fig. 6A, 6B, S5A and S6B). The mitigation of the number of peaks and hence of accessibility at the transcription starting sites (TSS) of these genes are demonstrated in Fig. 6C. Consistent with our hypothesis and the data displayed in earlier figures, blocking SUMOylation induces a switch from an open to a close chromatin state at promoters of genes that promote synovial sarcomagenesis (Fig. 6D).

**Figure 6.**
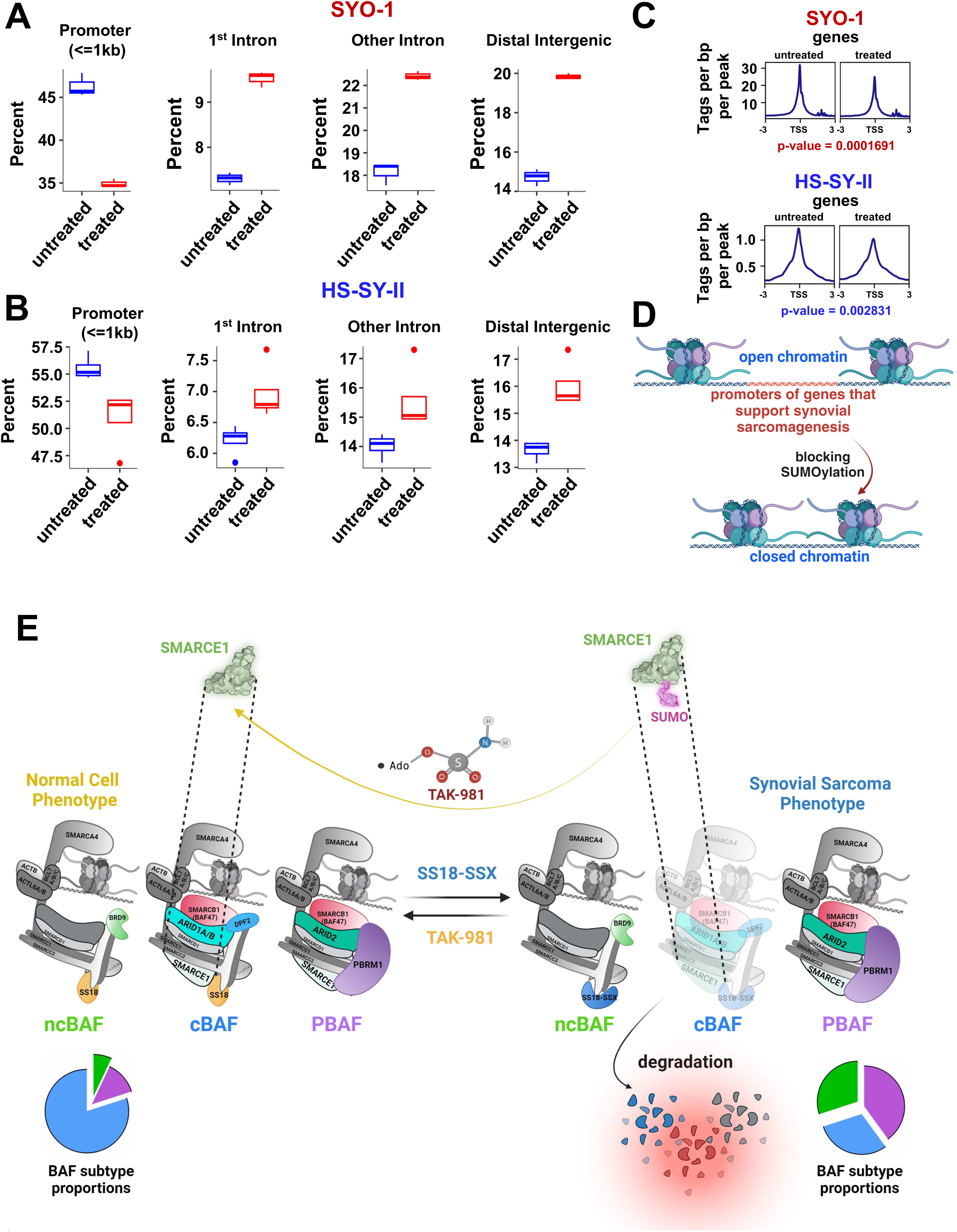
**A) (A) and B)** Boxplots demonstrating distribution of SYO-1 and HS-SY-II Atac seq peaks genome-wide, focusing on downregulated RNA-Seq normalized genes for HS-SY-II transduced with SS18-SSX shRNA ((filtered for DEGs with log2 (fold change) > +1). **C)** SYO-1 and HS-SY-II Atac seq profiles for downregulated RNA-Seq normalized genes of HS-SY-II cells transduced with SS18-SSX shRNA (the same list of downregulated genes used in Figure 6A and 6B), centered on TSS (transcription starting sites). **D)** Schema of changes at the chromatin following TAK-981 treatment in SS. **E)** Model of TAK-981 efficacy. TAK-981 deSUMOylates SMARCE1, leading to loss of RNF4-mediated degradation. SMARCE1 stabilization leads to cBAF complex stabilization at chromatin, and relative loss of ncBAF at chromatin. The result is inhibition of the SS18-SSX-ncBAF-driven transcriptome.

### TAK-981 has activity in synovial sarcoma *in vitro* and *in vivo* by inducing DNA damage

We next attempted to elucidate the mechanism that leads to SS toxicity after treatment with TAK-981. Consequently, we conducted gene set enrichment analysis (Subramanian et al., 2005) of the ATAC-seq data of the SYO-1 and HS-SY-II treated cells and realized that several among the top 20 downregulated pathways intersected on DNA damage (e.g. ‘’DNA repair’’, ‘’cell cycle’’, ‘’mismatch repair’’, ‘’base excision repair’’ etc. were suppressed) (Fig. S7A, S7B). Similar were the pathways obtained from the RNA-seq data from the experiments performed in the same cell lines (Fig. 4A). We also confirmed these data at the protein level with a well characterized DNA damage marker (p-H2AX (Ser139), also known as γ-H2AX (Rogakou et al., 1998)) (Fig. S7F).

Seeking a direct link between the observed rebalancing of the BAF complexes and the induction of DNA damage, we focused on the promoters of genes associated with reduced SS18:SSX (in HS-SY-II and SYO-1 cells) and SMARCA4 (in HS-SY-II ChIP seq peaks. In these analyses, we identified motifs of several TFs (transcription factors) of DNA repair related genes (Fig. S7C, S7D and S7E) pointing to a direct correlation between the binding of the BAF complexes and the attenuation of the DNA damage response.

We next tested TAK-981 *in vivo*, as dosed previously (Langston *et al*., 2021), in two synovial sarcoma cell line xenograft models (HS-SY-II and SYO-1) and one PDX model (SS.PDX). All tumors were grown in NOD Scid II2r Gamma (NSG) mice. The HS-SY-II and SYO-1 models were treated with 25 and 50 mg/kg of TAK-981 via tail vein injection, three consecutive days a week, or with no drug. The HS-SY-II tumors demonstrated tumor growth inhibition at both doses (Fig. 7A and Fig. S8A, S8B) while the SYO-1 tumors displayed tumor shrinkage at the low dose and were not detectable by calipers at the high dose of 50 mg/kg (Fig. 7B and Fig. S8A, S8B). The enhanced sensitivity of SYO-1 tumors is consistent with the more marked rebalancing of cBAF at the chromatin (Fig. 3D). Furthermore, to explore the lowest efficient dose of the drug *in vivo*, we treated a PDX model with 7.5 mg/kg of TAK-981, for three consecutive days a week, for two cycles (days of treatment: 1,2,3,8,9,10) and monitored the tumor growth after the 10^th^ day of the study, for a total period of four weeks. Even at these doses, tumor growth was almost completely blocked (Fig. 7C and Fig. S8A, S8B). Representative tumors of the SYO-1 model, before and after drug treatment, were subsequently excised and stained by immunohistochemistry for cleaved caspase-3 and γ-H2AX to corroborate cell death and the DNA damage induction noted in our *in vitro* experiments (Fig. 7D and S8C), as well as for SMARCE1, SMARCB1 (BAF47) and ARID1A expression, verifying the elevation of the specific fellow cBAF complex components (Fig. 7D and S8C), consistent with our *in vitro* studies. The staining was also quantified and the H-score for the expression of each protein was obtained, verifying the observed differences (Fig. 7E). There was no weight loss of the mice for any of our *in vivo* treatments, indicating that the administration of TAK-981, at the doses tested, is well tolerated (Fig. S8D, S8E and S8F), consistent with the emerging clinical data.

**Figure 7.**
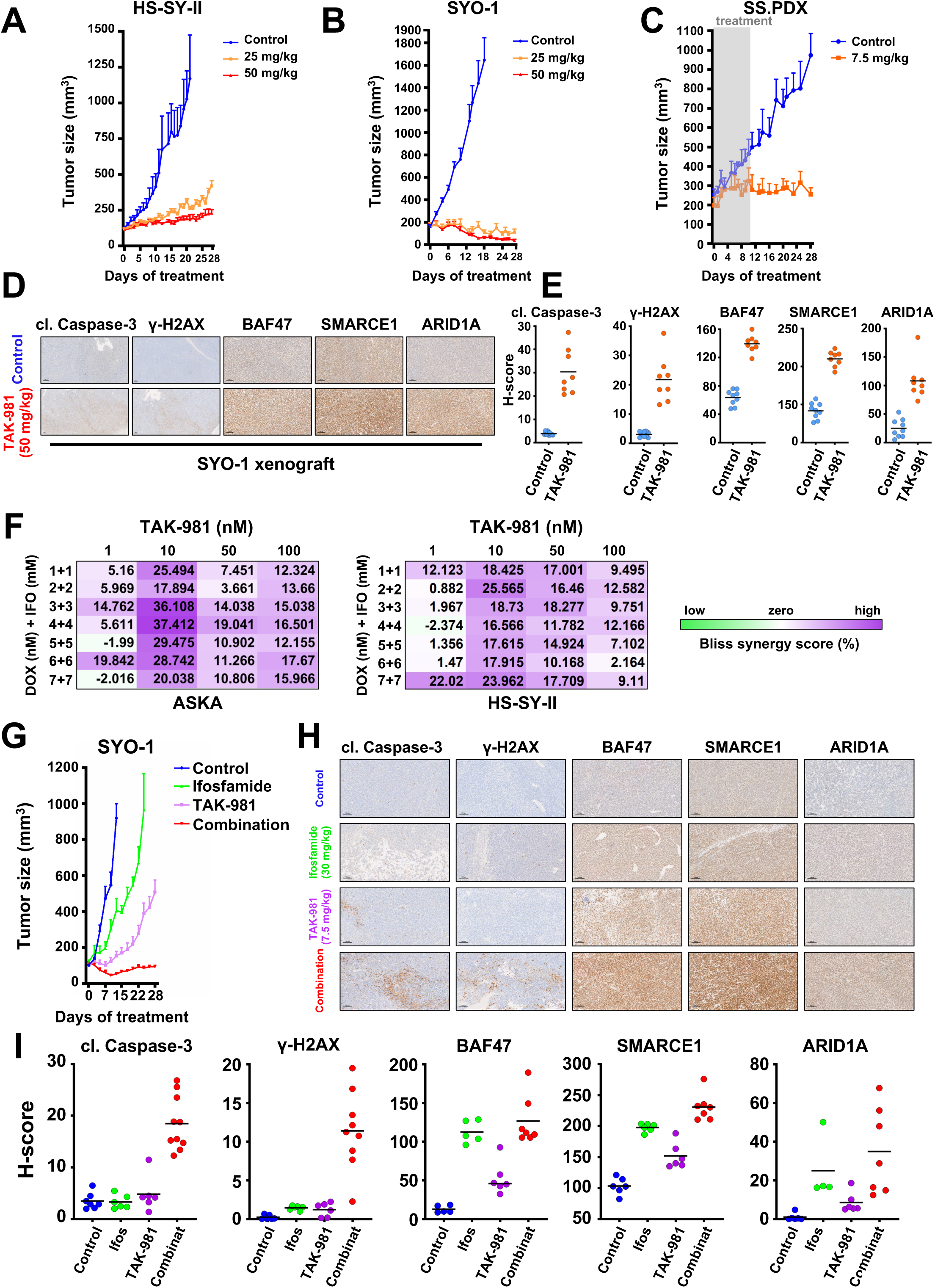
**A)** HS-SY-II tumor-bearing NSG mice were treated with 25mg/kg or 50mg/kg TAK-981 three consecutive days a week (Tuesday-Thursday) or control (no treatment) and tumors were monitored by caliper at least 3/w (n=6 for control, 5 for 25 mg/kg and 6 for 50 mg/kg). **B)** SYO-1 tumor-bearing NSG mice were treated with 25mg/kg or 50mg/kg TAK-981 3/w or control (no treatment) and tumors were monitored by caliper at least 3/w (n=8 for control, 8 for 25 mg/kg and 9 for 50 mg/kg). **C)** SS.PDX tumor-bearing NSG mice were treated with 7.5mg/kg TAK-981 or control three consecutive days a week (Tuesday-Thursday) the first and second week of the experiment. The drug administration was paused after the second round of treatment (day 10) and the mice were monitored for tumor growth for a total period of 28 days (n=6 for control, 5 for 7.5 mg/kg). **D)** Representative images of IHC analysis of the indicated antibodies in SYO-1 tumors from experiment shown in (B). Scale bars = 100 μm. **E)** H-scores of staining from (D). **F)** Bliss Sum synergy scores were obtained following treatment with the indicated drugs at the indicated concentrations. **G)** SYO-1 tumor-bearing NSG mice (n=x per cohort) were treated with TAK-981 via tail vein injection at a dosage of 7.5 mg/kg, three consecutive days a week (Tuesday-Thursday) for four weeks (28 days) and ifosfamide via intraperitoneal injection at a dosage of 30 mg/kg, for three consecutive days a week (Tuesday-Thursday) the first and fourth week of the experiment. Tumor measurements were performed every other day by calipers, and the average tumor volume + SEM for each cohort is displayed (n=7 for control, 9 for ifosfamide, 8 for TAK-981 and 8 for combination treatment). **H)** Representative images of IHC analysis of the indicated antibodies in SYO-1 tumors from the experiment shown in (G). Scale bars = 100 μm. **I)** H-scores of staining from (H).

### TAK-981 sensitizes SS to cytotoxic chemotherapy *in vitro* and *in vivo*

Doxorubicin (DOX)-based chemotherapy remains the commonly applied first-line therapy for SS, with the addition of ifosfamide (IFO) providing some benefit (Judson et al., 2014). Since both DOX and IFO are DNA damaging chemicals, DOX by intercalation and IFO by alkylation (Hartley et al., 1999; Willits et al., 2005; Yang et al., 2014), we hypothesized that the addition of TAK-981 would synergize with these by inducing enhanced DNA damage. We found significant synergy of DOX/IFO with TAK-981 (72h) using a Bliss-Sum matrix and calculating a percentage over the Bliss score ^39^ across multiple doses of the drugs (Fig. 7F and S9A) (Alkhatib et al., 2019; Morgan and Cranmer, 2014). In line with our hypothesis, fragmentation of DNA was more pronounced in combination treated cells than following DOX plus IFO or TAK-981 alone, as assessed by the COMET assay (Olive and Banáth, 2006) (portrayed by the ‘’tails’’ escorting the stained nuclei) (Fig. S9B). These data provide a rationale for combining TAK-981 with DOX and/or IFO.

To minimize the adverse effects from the usage of multiple agents, we attempted to test the efficacy of the combination of TAK-981 with IFO *in vivo*, without the presence of DOX. IFO alone has been used several times for the therapy of soft tissue sarcoma patients (Benjamin et al., 1993; Blay et al., 2023; Lee et al., 2011; Rosen et al., 1994). and there are cases where DOX and IFO together haven’t shown greater activity than IFO alone (Desar et al., 2018). The administration of IFO+TAK-981 (depicted in Fig. S9C) demonstrated impressive combination activity compared to single-agent activity, with the tumors treated with the combination shrinking (Fig. 7G, S9D and S9E). Noteworthy, no weight loss was detected in the combination-treated mice, suggesting tolerability (Fig. S9F). Finally, IHC analysis of the tissues and its quantification by calculating the H-score of the combination cohort revealed enhanced staining for cleaved caspase-3, γ-H2AX, and for the three cBAF (ARID1A) and cBAF/PBAF (BAF47, SMARCE1) - specific components compared to the tumors of the TAK-981 or the IFO treatment alone (Fig. 7H, 7I and S9G).

### TAK-981 blocks tumor growth in a mouse model with conditional SS18::SSX expression

To further test the effect of TAK-981 *in vivo* we utilized a genetic SS mouse model that conditionally expresses SS18::SSX2 with littermate-controlled cohorts of *hSS2* mice (homozygous for *hSS2* at the *Rosa26* locus) (Fig. 8A). TATCre was injected to initiate SS tumors in the *hSS2* mice (Fig. 8A), followed by 25 mg/kg of TAK-981 or vehicle administration. Consistent with the mouse models harboring human SS tumors, TAK-981 treatment led to significantly reduced tumors compared to the vehicle-treated tumors (Fig. 8B, 8C and 8D). RNA-seq analysis of vehicle-treated tumors and TAK-981 treated tumors demonstrated clustering of gene expression between the two cohorts (Fig. 8E) and pathways associated with DNA repair and the cell cycle were significantly downregulated in TAK-981 treated tumors (Fig. 8F, S10A and S10B). In contrast, pathways related to muscle function, including those involved in cardiac muscle contraction and hypertrophic cardiomyopathy, as well as to ECM receptor interaction and focal adhesion, exhibited a marked upregulation in TAK-981 treated tumors (Fig. 8F, S10A and S10C). These data demonstrate that SUMOylation pathway inhibition in a conditionally expressed SS18::SSX2 mouse model leads to a reduction of the neuronal characteristics of synovial sarcoma tumors and a concurrent gain of muscle-related phenotypes, as was also previously demonstrated by the RNA-seq, ChIP-seq and ATAC-seq analyses of our *in vitro* experiments (Fig. 4A, 5A, 5D, 6E and 6F). Additionally, a significant overlap was noticed between the up- and down-regulated genes of our mouse model and the 100 most up- and down-regulated genes after SSX knock down (Fig. 8G and 8H). TAK-981 induced cell death was corroborated by western blotting (Fig. 8I), as well as the elevation in the expression of BAF47 and SMARCE1 (Fig. 8J, 8K and S10D), supporting restoration of cBAF complexes across our *in vitro* and *in vivo* models in response to the inhibition of SUMOylation in SS.

**Figure 8.**
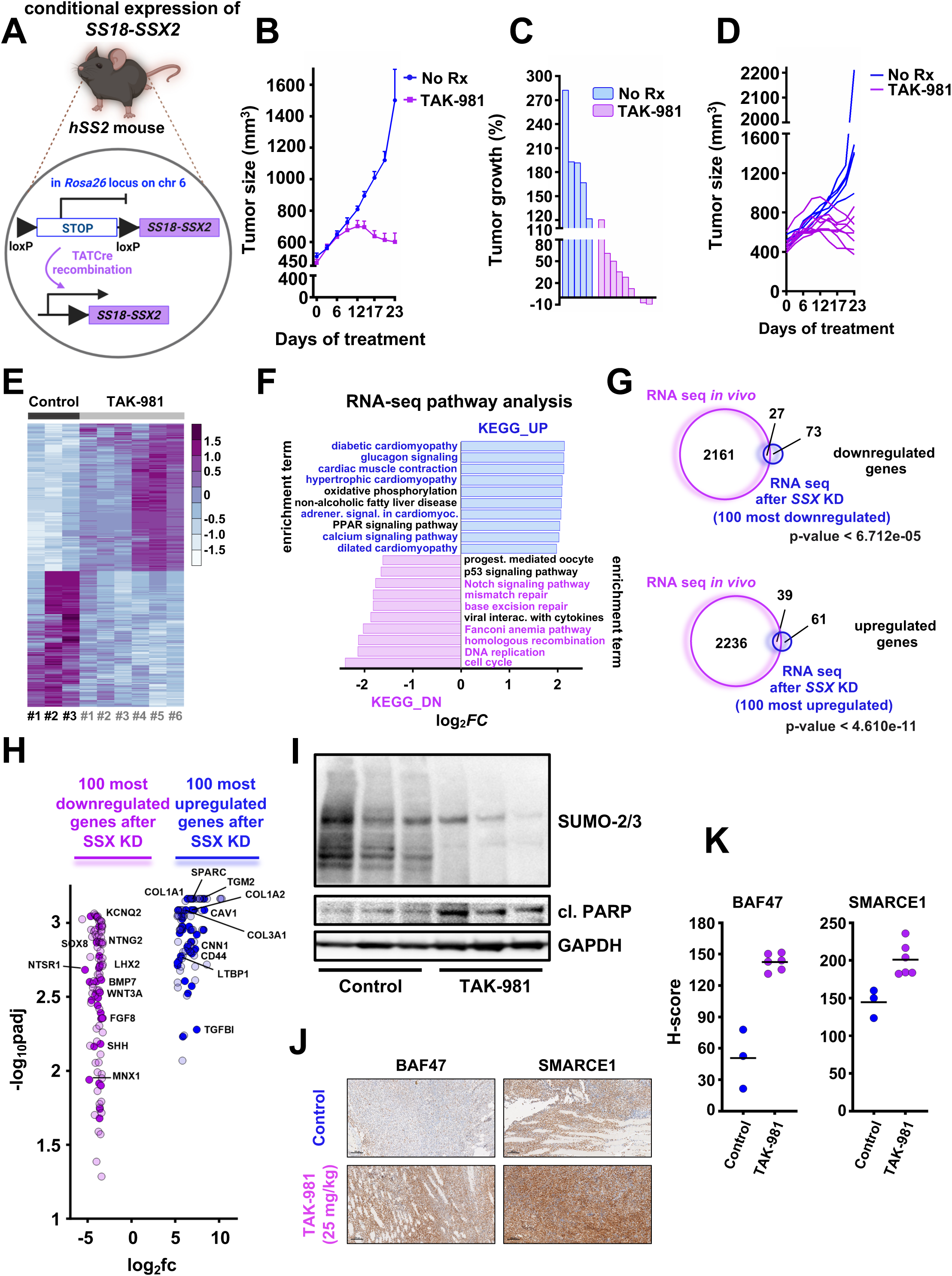
**A)** Schema of the design of the mouse experiment. **B)** hSS2 mice were treated with 25mg/kg 3/w or vehicle (control) and tumors were monitored by caliper at least 3/w (n=5 for control and n=9 for the TAK-981 treatment cohort) **C)** Waterfall plot of data from (B). **D)** Individual tumor growth from the experiment shown in (B and C). **E)** Heatmap of the RNA expression profiles of the 1000 most differentiated genes following TAK-981 treatment, after RNA seq analysis in the indicated hSS2 tumors (n=3 for control and n=6 for TAK-981 treatment). **F)** Pathway analysis of RNA-seq data from hSS2 tumors. **G)** Venn diagrams of significant gene changes caused by TAK-981 in hSS2 mice and the 100 most up- or down-regulated genes *in vitro* following SS18-SSX knockdown in HS-SY-II cells. **H)** Volcano plot of the 100 most down- or upregulated genes following SS18-SSX knockdown *in vitro* plotted with the common downregulated genes (purple dots) or common upregulated genes (blue dots) after RNA seq in the hSS2 tumors. **I)** Representative tumors from the control and TAK-981 treated cohort were harvested 2-3h after the last TAK-981 administration and tumor lysates (supplemented with 0.5M of N-ethylmaleimide (NEM)) were subjected to western blot analyses and probed for the indicated proteins. **J)** Representative images of IHC analysis of the indicated antibodies in the tumors from the experiment shown in (B). Scale bars = 100 μm. **K)** H-scores of staining from (J).

## Discussion

SS18::SSX is a fusion oncogene found invariably in SS tumors that assembles into BAF complexes and leads to the eviction of the wildtype SS18 protein from the complex^13^. SS18::SSX exerts its oncogenic effect by rebalancing BAF complexes from a cBAF dominated landscape to one with increasing relative prevalence of ncBAF, and to a lesser extent, PBAF. As such, reversing this re-balancing is an attractive therapeutic strategy for SS. Indeed, after either ncBAF complex activity or increasing cBAF complex activity has anti-SS effects^10-12^. In this study, we demonstrate that this rebalancing can be achieved through blocking SUMOylation with the small molecule inhibitor, TAK-981.

Overall, SUMOylation appears to most prominently impact chromatin structure and transcriptional function(Boulanger et al., 2021; Kim et al., 2021; Roy and Sadanandom, 2021; Wu et al., 2020). SUMOylation often negatively affects protein expression, but it can also increase protein expression: For instance, since SUMOylation and ubiquitination compete for lysines, SUMOylation can lead to hinderance of ubiquitination and thus stabilization of the effected proteins (Flotho and Melchior, 2013; Park et al., 2007). Conversely, the SUMO-dependent E3 ubiquitin ligase, RNF4, can degrade SUMOylated proteins leading to the loss of their expression(Han et al., 2021; Keiten-Schmitz et al., 2019). Indeed, in our study we found that SUMO-dependent SMARCE1 depletion was RNF4 co-dependent. Consistent with recent findings in CCM, where SMARCE1 is a stabilizer of cBAF complexes on chromatin (St Pierre *et al*., 2022), we found increased SMARCE1 presence was sufficient to stabilize cBAF complexes. This was sufficient to shift BAF complexes away from the SS state, with marked loss of the ncBAF-driven SS signature. Overall, TAK-981 acts as a surrogate SS18::SSX inhibitor in SS and has substantial anti-SS activity *in vitro* and *in vivo*. Of note, consistent with increased SMARCE1 expression resulting in toxicity in SS, our attempts to reconstitute wt SMARCE1 and SMARCE1 mutated at the sites of SUMOylation (K92 and K277) either constitutively or through an induced system, resulted in toxicity.

Preclinical studies have also demonstrated a role for TAK-981 in immune cells, changes that may promote cancer treatment benefit (Lam et al., 2023; Nakamura et al., 2022). This study did not evaluate a role of the immune system in SS, and further studies should be performed to elicit any additional effects TAK-981 may have in this context.

Furthermore, while our data suggests TAK-981 disruption of several motifs involved in DNA repair are downregulated following TAK-981 treatment, which may be the direct link to the DNA damage seen following BAF complex redistribution. While our data indicate that this is at least in part due to the rebalancing of BAF complexes, SUMOylation plays a direct role in the DNA damage response through other targets of SUMOylation (Claessens et al., 2023; Gasser and Stutz, 2023). Thus, further studies are needed to better characterize the mechanism of DNA damage following TAK-981 treatment in SS. Metastatic SS continues to have a poor prognosis due to poor disease control by available therapies. DOX-based chemotherapy remains the commonly practiced standard therapy in many centers, and adding IFO to DOX increases overall response rates and progression-free survival, but neither significantly extend overall survival (Edmonson et al., 2003; Judson *et al*., 2014). Pazopanib is a pan-kinase inhibitor approved for SS but has a low response rate (Kojima et al., 2022; Yoo et al., 2015). Merck’s PD-1 inhibitor, Pembrolizumab, was recently evaluated in soft tissue sarcomas including 10 SS cases with only one patient having a response (Tawbi et al., 2017). In addition, BRD9 degraders have been tested clinically, however the most advanced (CFT8364; C4 Therapeutics) has recently been discontinued. In a phase I trial across multiple patients with metastatic solid tumors or refractory/relapsed lymphoma, TAK-981 demonstrated tolerability and a RP2D of 90mg BIW was established. TAK-981 is also currently being evaluated in combination with pembrolizumab (NCT04381650). Of note, to our knowledge, TAK-981 has not been evaluated in any SS patients. Overall, we demonstrate a relative increased expression and activation of cBAF results in loss of the ncBAF-driven SS signature, leading to toxicity in SS, and that this can be accomplished with the in-clinic SUMOylation inhibitor, TAK-981. Our data provides rationale to test TAK-981 in SS patients, either alone or combined with cytotoxic chemotherapy.

## ACKNOWLEDGEMENTS

We thank Nicolo Riggi, Gaylor Boulay, Adam Crystal and Ronald Hay for invaluable support, feedback and conversation. This work was supported by the National Institutes of Health/National Cancer Institute (NIH/NCI) grant number 1R01CA272710-01A1 (to A.C.F.), a Department of Defense Rare Cancer Research Program grant, award number HT9425-23-1-1017 (to A.C.F.), and a Department of Defense Rare Cancer Research Program grant, award number W81XWH-22-1-0938 (to S.R.). A.B. received funding from the European Research Council (ERC) under the European Union’s Horizon 2020 research and innovation programme (grant agreement number 805338) and from the NIH/NCI U54CA231652. R.L. was supported by an NIH/NIAID grant (R01AI141410), Research Scholar Grant (134703-RSG-20–054-01-MPC) from the American Cancer Society and start-up funds from the University of Pittsburgh Medical Center (UPMC) Hillman Cancer Center. T.S. was supported by NIH grant R01 AI51051. Data was generated in the Genome Sequencing Facility, which is supported by UT Health San Antonio, NIH-NCI P30 CA054174 (Cancer Center at UT Health San Antonio), NIH Shared Instrument grant 1S10 OD030311-01 (S10 grant), and CPRIT Core Facility Award (RP220662). Z.L. is supported by NIH NCI R50 CA265339. Services and products in support of the current research project were also generated by the Virginia Commonwealth University Cancer Mouse Models Core Laboratory, and the Tissue and Data Acquisition and Analysis Core Laboratory, both supported, in part, with funding to the Massey Cancer Center from NIH-NCI Cancer Center Support Grant P30 CA016059.

## AUTHOR CONTRIBUTIONS

**Conceptualization**, K.V.F., and A.C.F.; **formal analysis**, K.V.F., C.K.F., J.L., K.Z., J.L.R., R.K., B.H., V.K., N.H., S.S., M.M.I., K.S.-F., L.L., A.S., K.M.D., A.J., E.I.A., Y.X., R.D.H., J.M.S., M.S., M.R.L., M.R.H., B.R.B., Z.L., S.A.B., A.M.S., J.P.L., M.H.M., K.V., R.L., A.B., A.P., J.E.K., T.S., M.G.D., K.B.J., S.K.R., A.C.F. ; **funding acquisition** K.B.J., S.K.R., T.S., A.C.F.; **investigation**, K.V.F., R.L., A.B., J.E.K., T.S., M.G.D., K.B.J., S.K.R., A.C.F.; **resources**, K.V.F., M.G.D., K.B.J., S.K.R., A.C.F.; **supervision**, A.C.F.; **writing – original draft**, K.V.F., A.C.F.; **writing – review and editing**, K.V.F., K.B.J., S.K.R., J.E.K., A.B., M.G.D., A.C.F.

## LEAD CONTACT STATEMENT

Further information and requests for resources and reagents should be directed to and will be fulfilled by the lead contact, Anthony Faber.

