## Supplemental material for "Targeting of SUMOylation leads to cBAF complex stabilization and disruption of the SS18::SSX transcriptome in Synovial Sarcoma"

#### Supplementary table

Table S1: PTMScan results

#### Supplementary figures

**Figure S1 (related to Fig. 1). A) - E)** DepMap (Dependency map) consortium genomic screen data in the four SS cell lines represented among the ~800 cancer cell lines. In this analyses from RNAi screens (Achilles + DRIVE + Marcotte), we noted sensitivity of siRNAs targeting several members of the SUMOylation pathway (*UBE2I* (*UBC9*), *SUMO2*, *PIAS1*, *UBA2* (*SAE2*) and *SAE1*), as analyzed by the Broad Institute scoring algorithm, DEMETER2. **B)** Schema of SUMOylation (created from Biorender).

**Figure S2 (related to Fig. 1). A)** Yamato SS cells were infected with a DOX-inducible control (shREN) or SS18-SSX knockdown (shSSX) and treated with 100nM TAK-981 for 36h and cell lysates (supplemented with 0.5M of N-ethylmaleimide (NEM)) were probed with the indicated antibodies. **B)** Crystal violet assays of SS cells untreated (No Rx) or treated with increasing concentrations of TAK-981 for five days. **C)** SS cells were treated with 100nM TAK-981 for 36h or left untreated (No Rx) and whole cell lysates were probed with the indicated antibodies.

**Figure S3 (related to Fig. 4). A)** Venn diagram of downregulated genes from RNA-seq following 100nM TAK-981 treatment for 36h in three SS cell lines. **B)** Venn diagram of upregulated genes from RNA-seq following 100nM TAK-981 treatment for 36h in three SS cell lines. **C) - E)** Pairwise comparisons between the three SS cell lines from RNA-seq following 100nM TAK-981 treatment for 36h. **F) - H)** Linear Regression analysis from pairwise comparisons between the three SS cell lines from RNA-seq following 100nM TAK-981 treatment for 36h. **I)** Plotted RNA-Seq fold changes of genes downregulated after TAK-981 treatment and pathway analysis. **J)** Quantitative RT-PCR validating gene expression results obtained by RNA-Seq for downregulated genes from (I). **K)** Plotted RNA-Seq fold changes of genes downregulated after TAK-981 treatment and pathway analysis. **L)** qPCR validation of genes from (K).

**Figure S4 (related to Fig. 5). A)** Venn diagrams of HS-SY-II cells treated with 100nM TAK-981 for 36h (and compared to untreated HS-SY-II cells) and significantly downregulated signals from ChIP-seq experiments (including SS18::SSX, SMARCA4, KDM2B and H3K27ac) and/or RNA-seq and/or the SS signature. **B)** The genes from the SS signature and those that overlap with reduced SS18::SSX ChIP-seq signals (orange boxes) or that do not (purple boxes) in HS-SY-II cells. **C)** Linear regression analysis of the correlation between SMARCA4 and SS18-SSX reduced ChIP-seq signals in HS-SY-II cells. **D)** Venn diagrams of HS-SY-II cells treated with 100nM TAK-981 for 36h (and compared to untreated HS-SY-II cells) and significantly upregulated signals from ChIP-seq experiments (including SS18::SSX, SMARCA4 and H3K27ac) and/or RNA-seq. **E)** Linear regression analysis of the correlation between SMARCA4 and SS18-SSX increased ChIP-seq signals in HS-SY-II cells. **F)** ChIP-seq profiles of H3K27ac and SMARCA4 (red = overlapping signals at commonly upregulated or downregulated genes) in HS-SY-II cells.

**Figure S5 (related to Fig. 5). A)** Venn diagram of SYO.1 cells treated with 100nM TAK-981 for 36h (and compared to untreated SYO-1 cells) and significantly downregulated signals from SS18-SSX ChIP-seq compared to the 100 most downregulated genes after SS18-SSX knockdown in HS-SY-II cells. **B)** Genes that overlap from (A) are plotted (orange dots indicate overlapping genes). **C)** Venn diagram of SYO-1 cells treated with 100nM TAK-981 for 36h (and compared to untreated SYO-1 cells) and significantly downregulated genes from RNA-seq data compared to significantly downregulated signals from SS18-SSX ChIP-seq experiments. **D)** Venn diagram of SYO-1 cells treated with 100nM TAK-981 for 36h (and compared to untreated SYO-1 cells) and significantly downregulated signals from SS18-SSX ChIP-seq experiments compared to significantly downregulated signals from KDM2B ChIP-seq experiments. **E)** Depth-normalized signal of SS18-SSX ChIP-seq and KDM2B ChIP-seq using IGV genome browser, reflecting losses in SS18-SSX and KDM2B peaks at the locus of genes supporting synovial sarcomagenesis in the SYO-1 cells. **F)** Venn diagram of SYO-1 cells treated with 100nM TAK-981 for 36h (and compared to untreated SYO-1 cells) and significantly downregulated genes compared to the SS signature. **G)** The genes from the SS signature and those that overlap with reduced SS18-SSX ChIP-seq signals (orange boxes) or that do not (yellow boxes) in SYO-1 cells. **H)** Venn diagram of SYO-1 cells treated with 100nM TAK-981 for 36h (and compared to untreated SYO-1 cells) and significantly downregulated signals from H3K27ac ChIP-seq experiments compared to significantly downregulated signals from SS18-SSX ChIP-seq experiments. **I)** Venn diagram of SYO-1 cells treated with 100nM TAK-981 for 36h (and compared to untreated SYO-1 cells) and significantly upregulated genes from RNA-seq data compared to significantly upregulated signals from SS18-SSX ChIP-seq experiments. **J)** Venn diagram of SYO.1 cells treated with 100nM TAK-981 for 36h (and compared to untreated SYO-1 cells) and significantly upregulated genes from H3K27ac ChIP-seq experiments compared to significantly downregulated signals from SS18-SSX ChIP-seq experiments. **K)** Volcano plot with the ChIP-seq profiles of H3K27ac and SS18-SSX (orange = overlapping reduced signals, red = overlapping increased signals at commonly upregulated or downregulated genes) in SYO-1 cells.

**Figure S6 (related to Fig. 6). A)** and **B)** Boxplots demonstrating distribution of the rest of the SYO.1 and HS-SY-II Atac seq peaks genome-wide (from Figure 6A and 6B), focusing on downregulated RNA-Seq normalized genes for HS-SY-II cells transduced with SS18-SSX shRNA (filtered like in Figure 6A and 6B). UTR, untranslated region.

**Figure S7 (related to Fig. 6). A)** and **B)** Gene set enrichment analysis using KEGG of downregulated and upregulated genes detected by Atac-seq, after treatment of SYO-1 and HS-SY-II cells with TAK-981 for 36h. **C)**, **D)** and **E)** Top 20 motifs enriched in the promoters of genes associated with downregulated peaks following TAK-981 treatment from the previous SS18-SSX ChIP seq data in SYO-1 (C) and HS-SY-II (D), as well as from the previous SMARCA4 ChIP seq data in HS-SY-II cells (E). **F)** Whole cell lysates from HS-SY-II, SYO-1 and SS.PDX synovial sarcoma cell lines, treated with 100 nM TAK-981 for 36h were prepared, subjected to immunoblotting, and probed for p-H2AX (Ser 139) and GAPDH.

**Figure S8 (related to Fig. 7). A)** Waterfall plots of HS-SY-II, SYO-1 or SS.PDX tumor-bearing NSG mice (from Fig. 7A, 7B and 7C) were treated with 25mg/kg, 50mg/kg TAK-981 (SYO-1 and HS-SY-II models) or 7.5 mg/kg (SS.PDX model) 3/w or control (no treatment) and tumors were monitored by caliper at least 3/w. **B)** Individual tumors were plotted from 7A, 7B, 7C. **C)** Representative images of IHC analysis of the indicated antibodies in SYO-1 tumors from the experiment shown in (7B). Scale bars = 100 µm. **D), E), F)** Weight changes from each of the three experiments from (7A, 7B, 7C).

**Figure S9 (related to Fig. 7). A)** Bliss Sum synergy scores were obtained following treatment for 72hrs with the indicated drugs at the indicated concentrations (like in Fig. 7F). **B)** Comet assay was performed following 36h of the indicated drug treatments (for ASKA and HS-SY-II cells) (combination is all three drugs). **C)** Schema of drug treatments **D)** Waterfall plot of SYO-1 tumor-bearing NSG mice from Fig. 7G. **E)** Individual tumor growth of tumor-bearing NSG mice from Fig. 7G. **F)** Weight changes from the in vivo experiment (Fig. 7G). **G)** Representative images of IHC analysis of the indicated antibodies in SYO-1 tumors from experiment shown in Fig. 7G and scored in Fig. 7I (like in Fig. 7H).

**Figure S10 (related to Fig. 8). A)** Volcano plot of the 500 most up- and the 500 most down-regulated genes after treatment with TAK-981 in the *in vivo* model (from Fig. 8). Some of the most characteristic up- or down-regulated genes are labeled **B)** Plotted RNA-Seq fold changes of genes downregulated after TAK-981 treatment (Fig. 8E-8H) and their associated pathways related to DNA repair, cell cycle and homeobox TFs. **C)** Plotted RNA-Seq fold changes of genes upregulated after TAK-981 treatment in the hSS2 mice (Fig. 8E-8H), and their associated pathways related to ECM receptor interaction, adrenergic signaling in cardiomyocytes, focal adhesion and protein digestion and absorption. **D)** Representative images of IHC analysis of the indicated antibodies in the mice from Fig. 8. Scale bars = 100 µm.

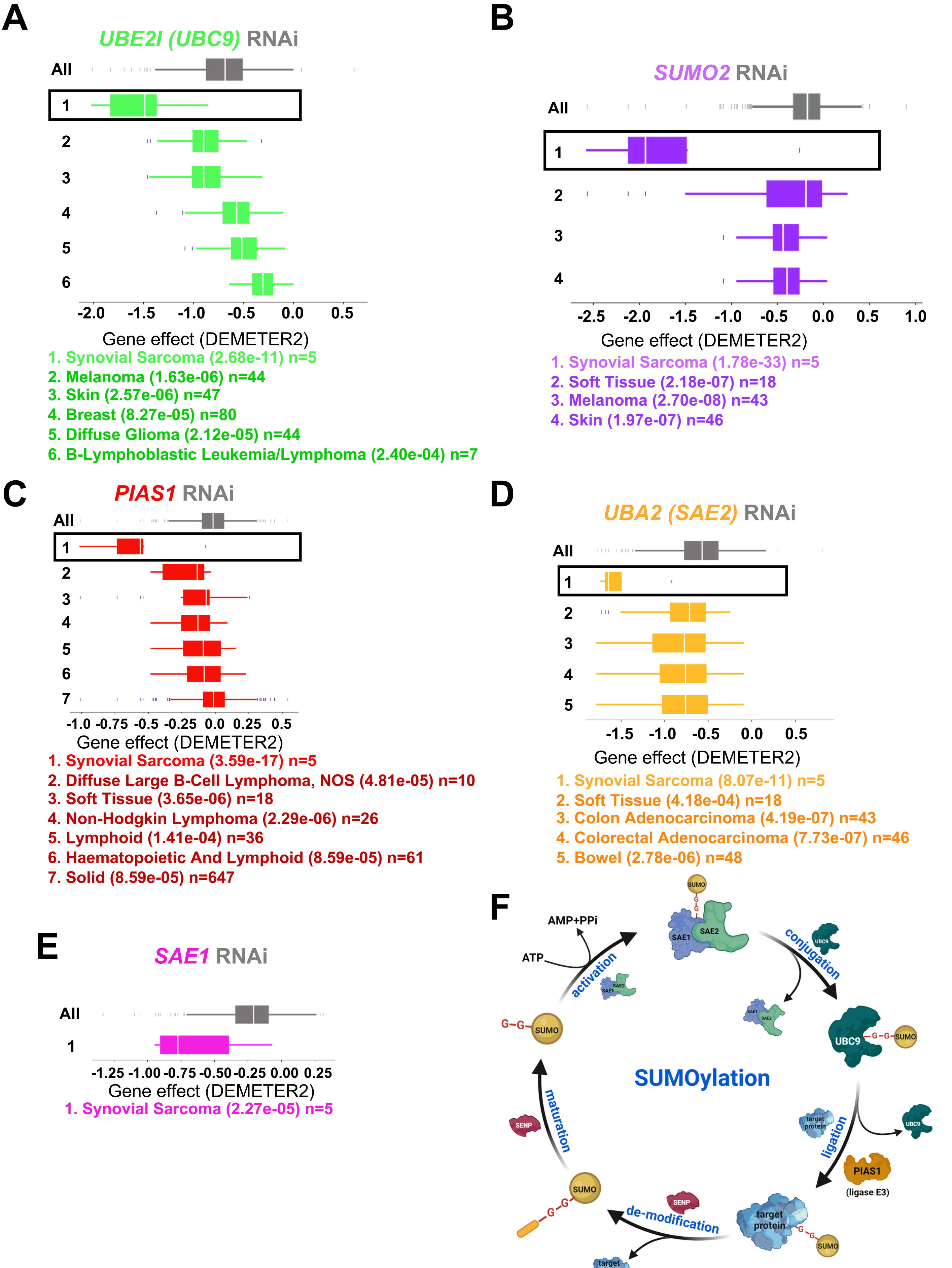
A
UBE2I (UBC9) RNAi
All
1
2
3
4
5
6
-2.0 -1.5 -1.0 -0.5 0.0 0.5
Gene effect (DEMETER2)
1. Synovial Sarcoma (2.68e-11) n=5
2. Melanoma (1.63e-06) n=44
3. Skin (2.57e-06) n=47
4. Breast (8.27e-05) n=80
5. Diffuse Glioma (2.12e-05) n=44
6. B-Lymphoblastic Leukemia/Lymphoma (2.40e-04) n=7
B
SUMO2 RNAi
All
1
2
3
4
-2.5 -2.0 -1.5 -1.0 -0.5 0.0 0.5 1.0
Gene effect (DEMETER2)
1. Synovial Sarcoma (1.78e-33) n=5
2. Soft Tissue (2.18e-07) n=18
3. Melanoma (2.70e-08) n=43
4. Skin (1.97e-07) n=46
C
PIAS1 RNAi
All
1
2
3
4
5
6
7
-1.0 -0.75 -0.5 -0.25 0.0 0.25 0.5
Gene effect (DEMETER2)
1. Synovial Sarcoma (3.59e-17) n=5
2. Diffuse Large B-Cell Lymphoma, NOS (4.81e-05) n=10
3. Soft Tissue (3.65e-06) n=18
4. Non-Hodgkin Lymphoma (2.29e-06) n=26
5. Lymphoid (1.41e-04) n=36
6. Haematopoietic And Lymphoid (8.59e-05) n=61
7. Solid (8.59e-05) n=647
D
UBA2 (SAE2) RNAi
All
1
2
3
4
5
-1.5 -1.0 -0.5 0.0 0.5
Gene effect (DEMETER2)
1. Synovial Sarcoma (8.07e-11) n=5
2. Soft Tissue (4.18e-04) n=18
3. Colon Adenocarcinoma (4.19e-07) n=43
4. Colorectal Adenocarcinoma (7.73e-07) n=46
5. Bowel (2.78e-06) n=48
E
SAE1 RNAi
All
1
-1.25 -1.00 -0.75 -0.50 -0.25 0.00 0.25
Gene effect (DEMETER2)
1. Synovial Sarcoma (2.27e-05) n=5
F
AMP+PPi
ATP
activation
SAE1
SAE2
SUMO
G-G
conjugation
UBC9
SUMOylation
maturation
SENP
ligation
target protein
PIAS1
(ligase E3)
de-modification


Figure S1

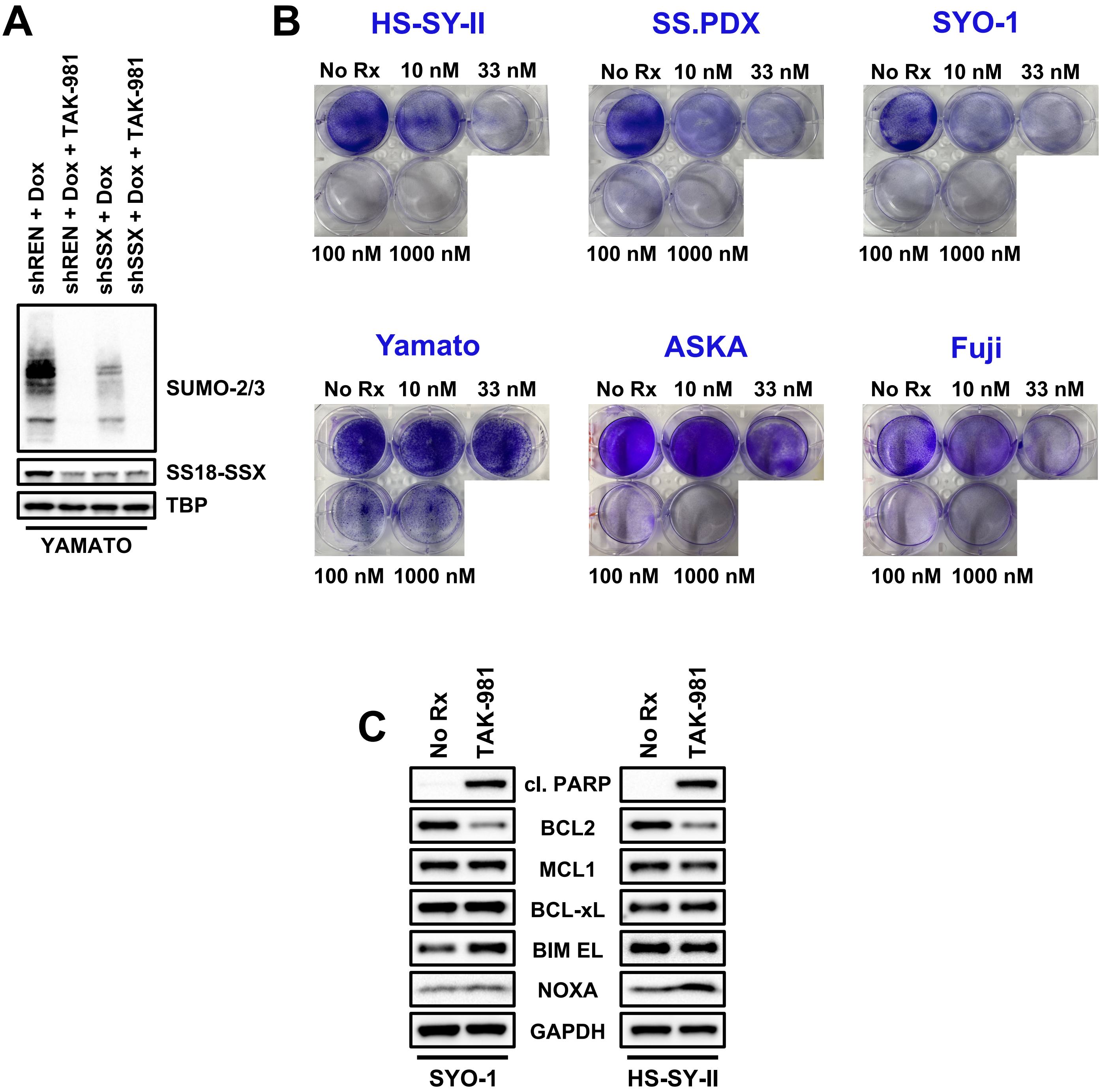
A
shREN + Dox
shREN + Dox + TAK-981
shSSX + Dox
shSSX + Dox + TAK-981
SUMO-2/3
SS18-SSX
TBP
YAMATO
B
HS-SY-II
No Rx 10 nM 33 nM
100 nM 1000 nM
SS.PDX
No Rx 10 nM 33 nM
100 nM 1000 nM
SYO-1
No Rx 10 nM 33 nM
100 nM 1000 nM
Yamato
No Rx 10 nM 33 nM
100 nM 1000 nM
ASKA
No Rx 10 nM 33 nM
100 nM 1000 nM
Fuji
No Rx 10 nM 33 nM
100 nM 1000 nM
C
No Rx
TAK-981
cl. PARP
BCL2
MCL1
BCL-xL
BIM EL
NOXA
GAPDH
SYO-1
HS-SY-II

Figure S2

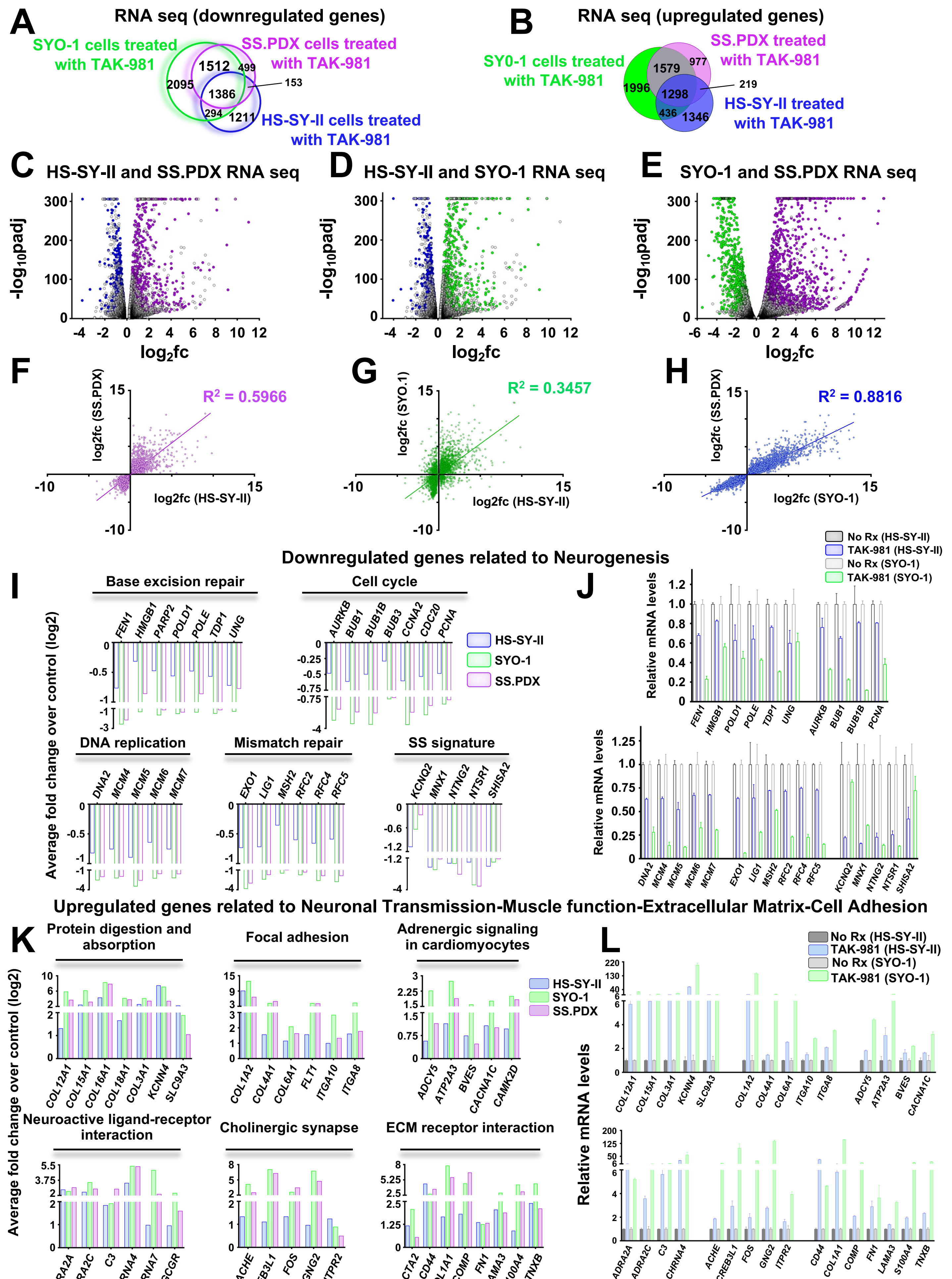
A
RNA seq (downregulated genes)
SYO-1 cells treated with TAK-981
SS.PDX cells treated with TAK-981
HS-SY-II cells treated with TAK-981
2095
1512
499
1386
153
294
1211
B
RNA seq (upregulated genes)
SY0-1 cells treated with TAK-981
SS.PDX treated with TAK-981
HS-SY-II treated with TAK-981
1996
1579
977
1298
219
436
1346
C
HS-SY-II and SS.PDX RNA seq
D
HS-SY-II and SYO-1 RNA seq
E
SYO-1 and SS.PDX RNA seq
-log10padj
log2fc
F
$R^2$ = 0.5966
log2fc (SS.PDX)
log2fc (HS-SY-II)
G
$R^2$ = 0.3457
log2fc (SYO.1)
log2fc (HS-SY-II)
H
$R^2$ = 0.8816
log2fc (SS.PDX)
log2fc (SYO-1)
Downregulated genes related to Neurogenesis
I
Base excision repair
FEN1
HMGB1
PARP2
POLD1
POLE
TDP1
UNG
Cell cycle
AURKB
BUB1
BUB1B
BUB3
CCNA2
CDC20
PCNA
HS-SY-II
SYO-1
SS.PDX
DNA replication
DNA2
MCM4
MCM5
MCM6
MCM7
Mismatch repair
EXO1
LIG1
MSH2
RFC2
RFC4
RFC5
SS signature
KCNQ2
MNX1
NTNG2
NTSR1
SHISA2
Average fold change over control (log2)
J
No Rx (HS-SY-II)
TAK-981 (HS-SY-II)
No Rx (SYO-1)
TAK-981 (SYO-1)
Relative mRNA levels
Upregulated genes related to Neuronal Transmission-Muscle function-Extracellular Matrix-Cell Adhesion
K
Protein digestion and absorption
COL12A1
COL15A1
COL16A1
COL18A1
COL3A1
KCNN4
SLC9A3
Focal adhesion
COL1A2
COL4A1
COL6A1
FLT1
ITGA10
ITGA8
Adrenergic signaling in cardiomyocytes
ADCY5
ATP2A3
BVES
CACNA1C
CAMK2D
HS-SY-II
SYO-1
SS.PDX
Neuroactive ligand-receptor interaction
ADRA2A
ADRA2C
C3
CHRNA4
CHRNA7
GCGR
Cholinergic synapse
ACHE
CREB3L1
FOS
GNG2
ITPR2
ECM receptor interaction
ACTA2
CD44
COL1A1
COMP
FN1
LAMA3
S100A4
TNXB
Average fold change over control (log2)
L
No Rx (HS-SY-II)
TAK-981 (HS-SY-II)
No Rx (SYO-1)
TAK-981 (SYO-1)
Relative mRNA levels

Figure S3

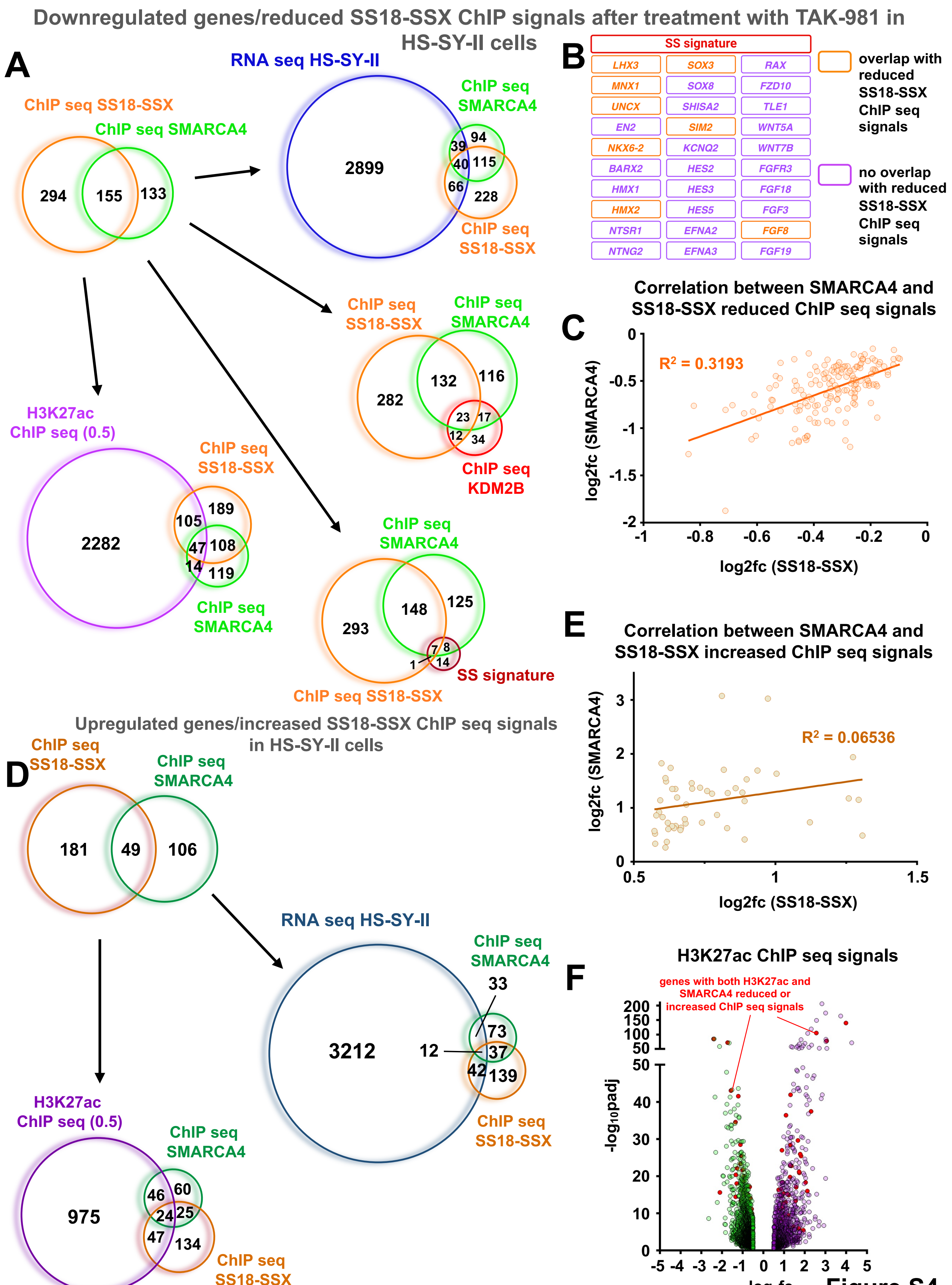
Downregulated genes/reduced SS18-SSX ChIP signals after treatment with TAK-981 in HS-SY-II cells
A
ChIP seq SS18-SSX
ChIP seq SMARCA4
294
155
133
RNA seq HS-SY-II
2899
ChIP seq SMARCA4
39
94
40
115
66
228
ChIP seq SS18-SSX
ChIP seq SS18-SSX
ChIP seq SMARCA4
282
132
116
23
17
12
34
ChIP seq KDM2B
H3K27ac ChIP seq (0.5)
2282
ChIP seq SS18-SSX
105
189
47
108
14
119
ChIP seq SMARCA4
ChIP seq SMARCA4
293
148
125
7
8
1
14
SS signature
ChIP seq SS18-SSX
B
SS signature
LHX3
SOX3
RAX
MNX1
SOX8
FZD10
UNCX
SHISA2
TLE1
EN2
SIM2
WNT5A
NKX6-2
KCNQ2
WNT7B
BARX2
HES2
FGFR3
HMX1
HES3
FGF18
HMX2
HES5
FGF3
NTSR1
EFNA2
FGF8
NTNG2
EFNA3
FGF19
overlap with reduced SS18-SSX ChIP seq signals
no overlap with reduced SS18-SSX ChIP seq signals
C
Correlation between SMARCA4 and SS18-SSX reduced ChIP seq signals
$R^2$ = 0.3193
log2fc (SMARCA4)
log2fc (SS18-SSX)
0
-0.5
-1
-1.5
-2
-1
-0.8
-0.6
-0.4
-0.2
0
Upregulated genes/increased SS18-SSX ChIP seq signals in HS-SY-II cells
D
ChIP seq SS18-SSX
ChIP seq SMARCA4
181
49
106
RNA seq HS-SY-II
3212
ChIP seq SMARCA4
33
73
12
37
42
139
ChIP seq SS18-SSX
H3K27ac ChIP seq (0.5)
975
ChIP seq SMARCA4
46
60
24
25
47
134
ChIP seq SS18-SSX
E
Correlation between SMARCA4 and SS18-SSX increased ChIP seq signals
$R^2$ = 0.06536
log2fc (SMARCA4)
log2fc (SS18-SSX)
3
2
1
0
0.5
1
1.5
F
H3K27ac ChIP seq signals
genes with both H3K27ac and SMARCA4 reduced or increased ChIP seq signals
$-log_{10}padj$
200
150
100
50
50
40
30
20
10
0
-5
-4
-3
-2
-1
0
1
2
3
4
5
$log_2fc$
Figure S4

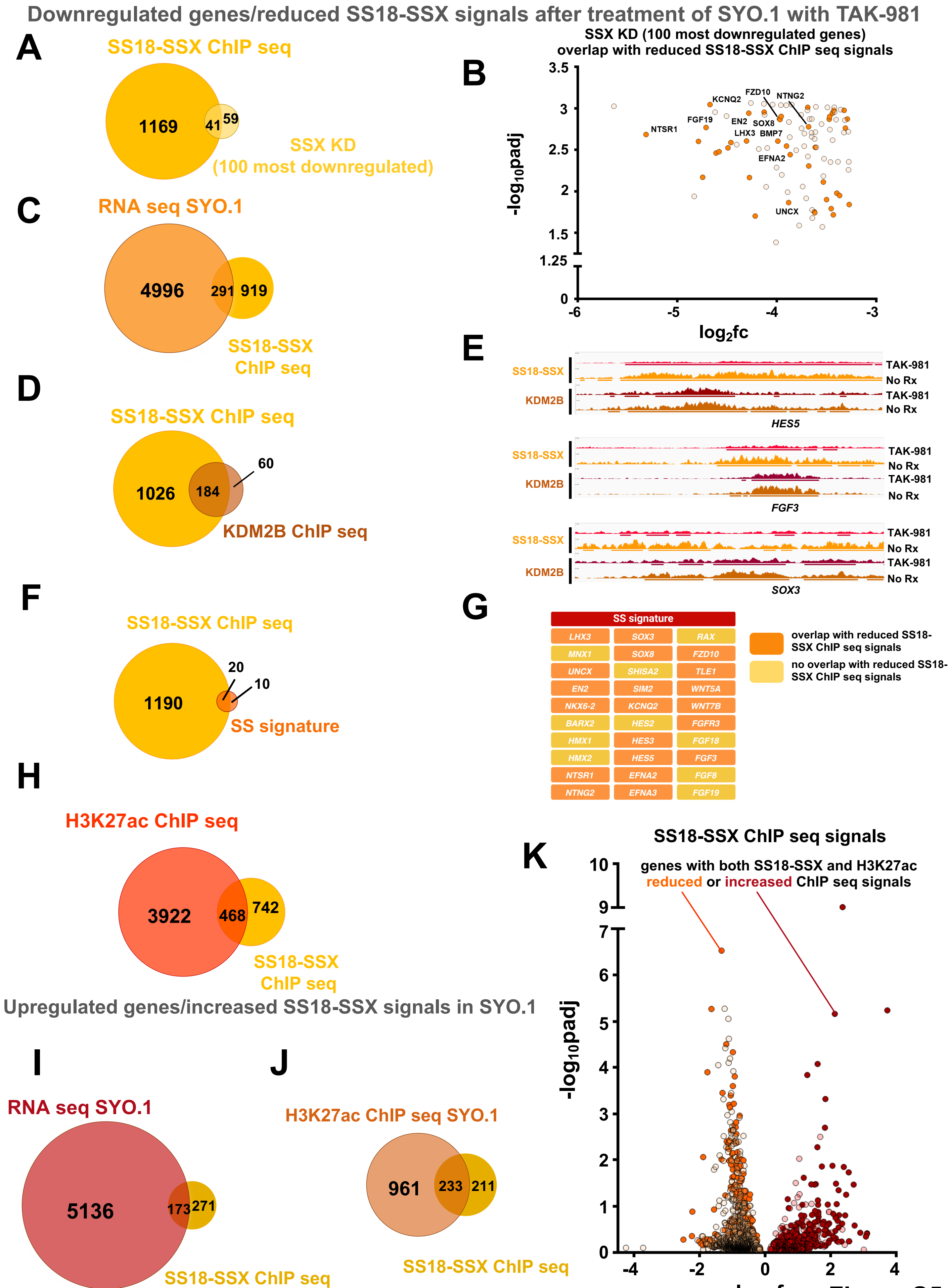
Downregulated genes/reduced SS18-SSX signals after treatment of SYO.1 with TAK-981
A
SS18-SSX ChIP seq
1169
41
59
SSX KD
(100 most downregulated)
B
SSX KD (100 most downregulated genes)
overlap with reduced SS18-SSX ChIP seq signals
-log10padj
log2fc
NTSR1
FGF19
KCNQ2
FZD10
NTNG2
EN2
SOX8
LHX3
BMP7
EFNA2
UNCX
C
RNA seq SYO.1
4996
291
919
SS18-SSX
ChIP seq
D
SS18-SSX ChIP seq
1026
184
60
KDM2B ChIP seq
E
SS18-SSX
KDM2B
TAK-981
No Rx
HES5
FGF3
SOX3
F
SS18-SSX ChIP seq
1190
20
10
SS signature
G
SS signature
LHX3
SOX3
RAX
MNX1
SOX8
FZD10
UNCX
SHISA2
TLE1
EN2
SIM2
WNT5A
NKX6-2
KCNQ2
WNT7B
BARX2
HES2
FGFR3
HMX1
HES3
FGF18
HMX2
HES5
FGF3
NTSR1
EFNA2
FGF8
NTNG2
EFNA3
FGF19
overlap with reduced SS18-SSX ChIP seq signals
no overlap with reduced SS18-SSX ChIP seq signals
H
H3K27ac ChIP seq
3922
468
742
SS18-SSX
ChIP seq
Upregulated genes/increased SS18-SSX signals in SYO.1
I
RNA seq SYO.1
5136
173
271
SS18-SSX ChIP seq
J
H3K27ac ChIP seq SYO.1
961
233
211
SS18-SSX ChIP seq
K
SS18-SSX ChIP seq signals
genes with both SS18-SSX and H3K27ac
reduced or increased ChIP seq signals
-log10padj
log2fc

Figure S5

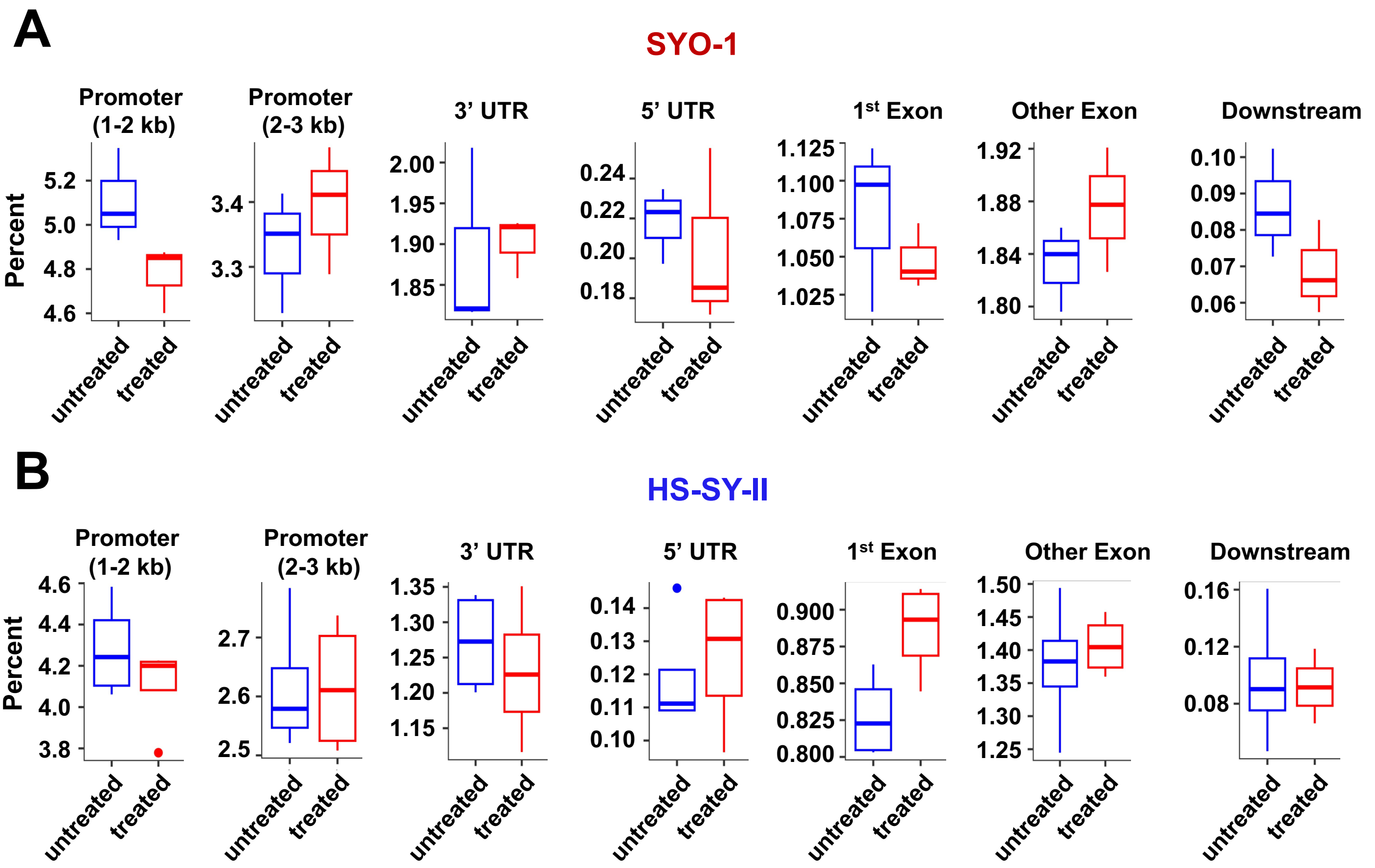
A
SYO-1
Promoter (1-2 kb)
Promoter (2-3 kb)
3’ UTR
5’ UTR
1st Exon
Other Exon
Downstream
Percent
untreated
treated
B
HS-SY-II
Promoter (1-2 kb)
Promoter (2-3 kb)
3’ UTR
5’ UTR
1st Exon
Other Exon
Downstream
Percent
untreated
treated

Figure S6

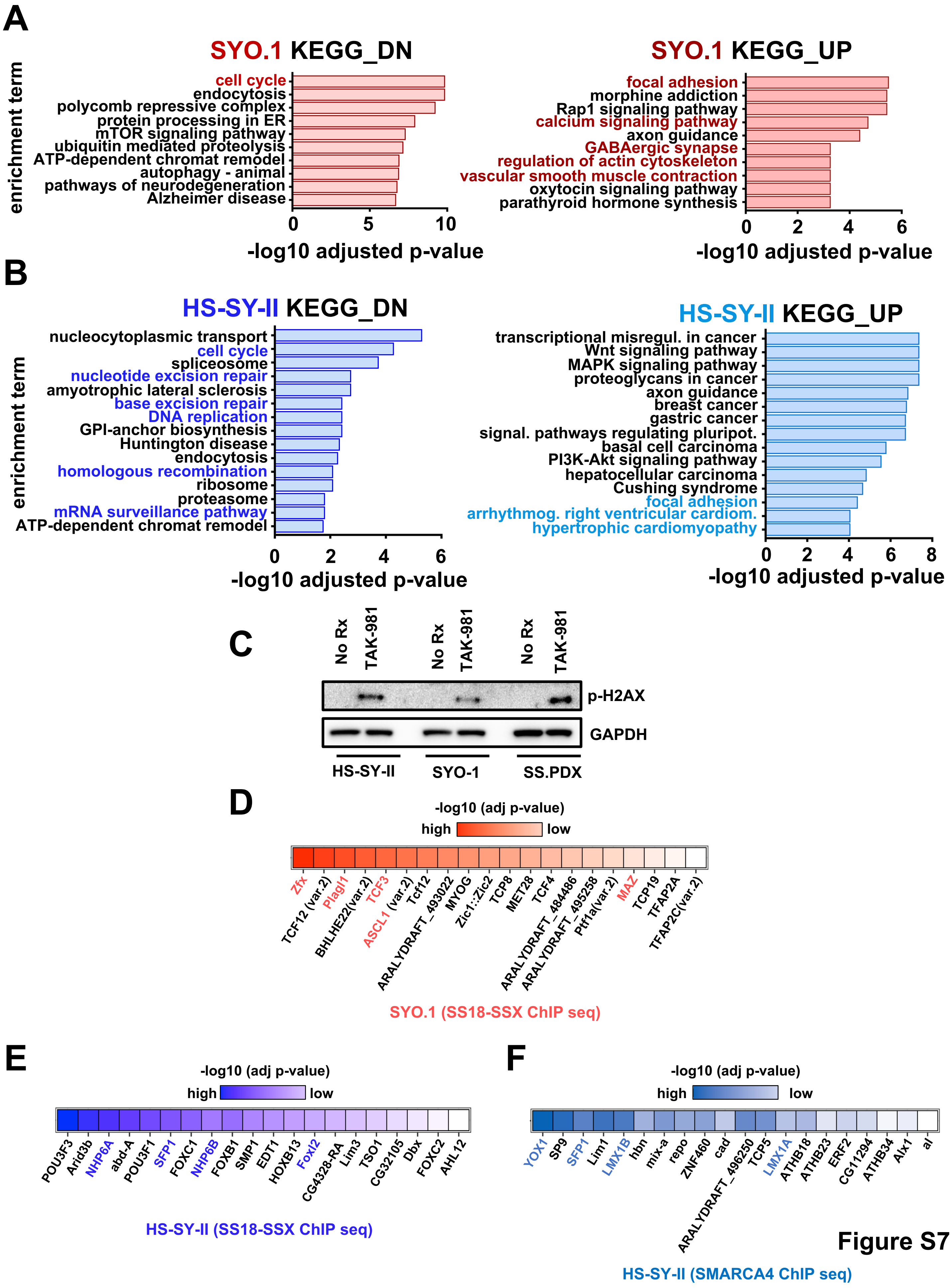
A
SYO.1 KEGG_DN
enrichment term
cell cycle
endocytosis
polycomb repressive complex
protein processing in ER
mTOR signaling pathway
ubiquitin mediated proteolysis
ATP-dependent chromat remodel
autophagy - animal
pathways of neurodegeneration
Alzheimer disease
0 5 10
-log10 adjusted p-value
SYO.1 KEGG_UP
focal adhesion
morphine addiction
Rap1 signaling pathway
calcium signaling pathway
axon guidance
GABAergic synapse
regulation of actin cytoskeleton
vascular smooth muscle contraction
oxytocin signaling pathway
parathyroid hormone synthesis
0 2 4 6
-log10 adjusted p-value
B
HS-SY-II KEGG_DN
enrichment term
nucleocytoplasmic transport
cell cycle
spliceosome
nucleotide excision repair
amyotrophic lateral sclerosis
base excision repair
DNA replication
GPI-anchor biosynthesis
Huntington disease
endocytosis
homologous recombination
ribosome
proteasome
mRNA surveillance pathway
ATP-dependent chromat remodel
0 2 4 6
-log10 adjusted p-value
HS-SY-II KEGG_UP
transcriptional misregul. in cancer
Wnt signaling pathway
MAPK signaling pathway
proteoglycans in cancer
axon guidance
breast cancer
gastric cancer
signal. pathways regulating pluripot.
basal cell carcinoma
PI3K-Akt signaling pathway
hepatocellular carcinoma
Cushing syndrome
focal adhesion
arrhythmog. right ventricular cardiom.
hypertrophic cardiomyopathy
0 2 4 6 8
-log10 adjusted p-value
C
No Rx
TAK-981
No Rx
TAK-981
No Rx
TAK-981
p-H2AX
GAPDH
HS-SY-II
SYO-1
SS.PDX
D
-log10 (adj p-value)
high
low
Zfx
TCF12 (var.2)
Plagl1
BHLHE22(var.2)
TCF3
ASCL1 (var.2)
Tcf12
ARALYDRAFT_493022
MYOG
Zic1::Zic2
TCP8
MET28
TCF4
ARALYDRAFT_484486
ARALYDRAFT_495258
Ptf1a(var.2)
MAZ
TCP19
TFAP2A
TFAP2C(var.2)
SYO.1 (SS18-SSX ChIP seq)
E
-log10 (adj p-value)
high
low
POU3F3
Arid3b
NHP6A
abd-A
POU3F1
SFP1
FOXC1
NHP6B
FOXB1
SMP1
EDT1
HOXB13
FoxI2
CG4328-RA
Lim3
TSO1
CG32105
Dbx
FOXC2
AHL12
HS-SY-II (SS18-SSX ChIP seq)
F
-log10 (adj p-value)
high
low
YOX1
SP9
SFP1
Lim1
LMX1B
hbn
mix-a
repo
ZNF460
cad
ARALYDRAFT_496250
TCP5
LMX1A
ATHB18
ATHB23
ERF2
CG11294
ATHB34
Alx1
al
HS-SY-II (SMARCA4 ChIP seq)

Figure S7

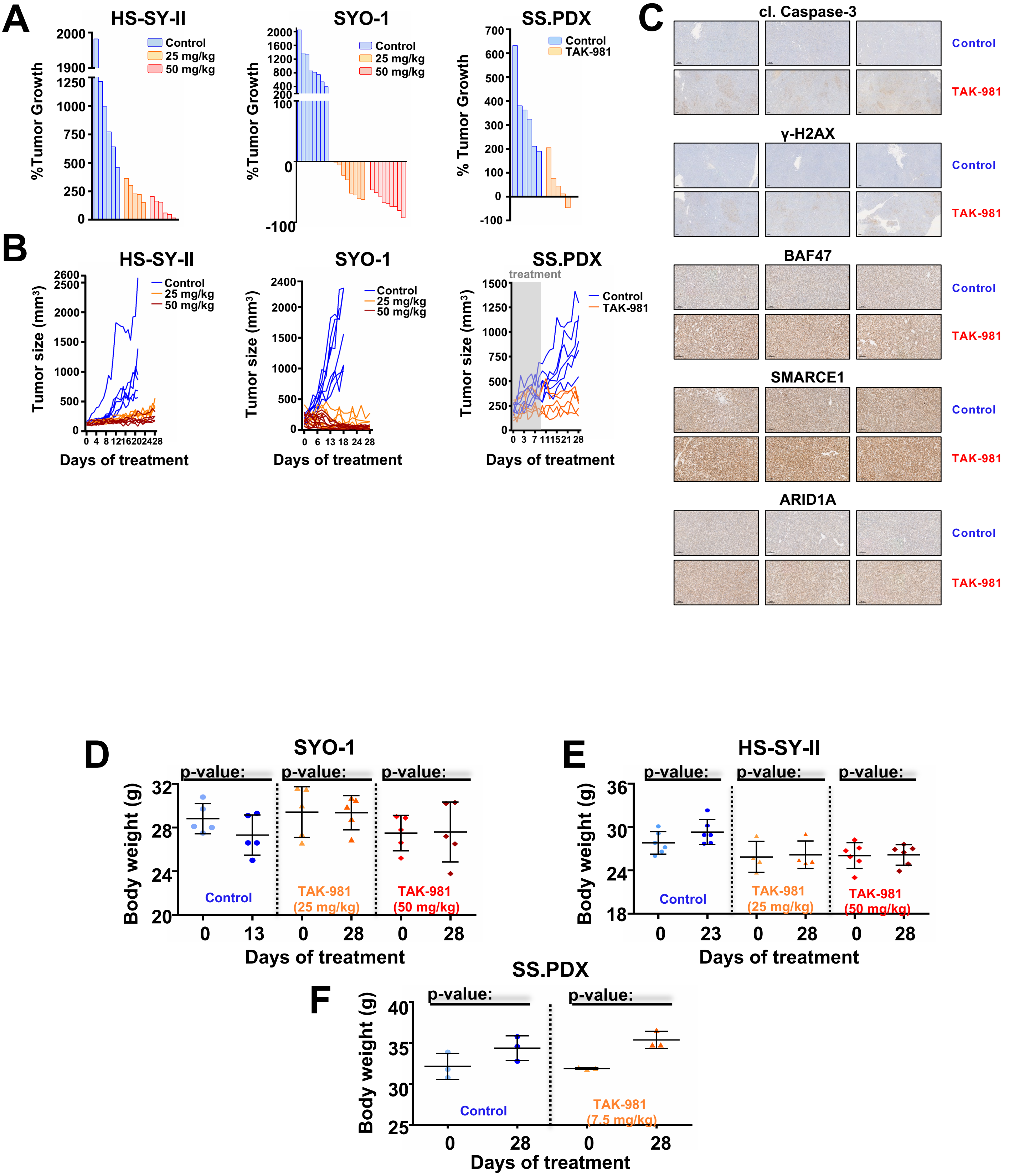
A
HS-SY-II
%Tumor Growth
Control
25 mg/kg
50 mg/kg
SYO-1
SS.PDX
% Tumor Growth
TAK-981
B
Tumor size (mm³)
Days of treatment
treatment
C
cl. Caspase-3
γ-H2AX
BAF47
SMARCE1
ARID1A
Control
TAK-981
D
SYO-1
E
HS-SY-II
F
SS.PDX
p-value:
Body weight (g)
TAK-981 (25 mg/kg)
TAK-981 (50 mg/kg)
TAK-981 (7.5 mg/kg)
Days of treatment

Figure S8

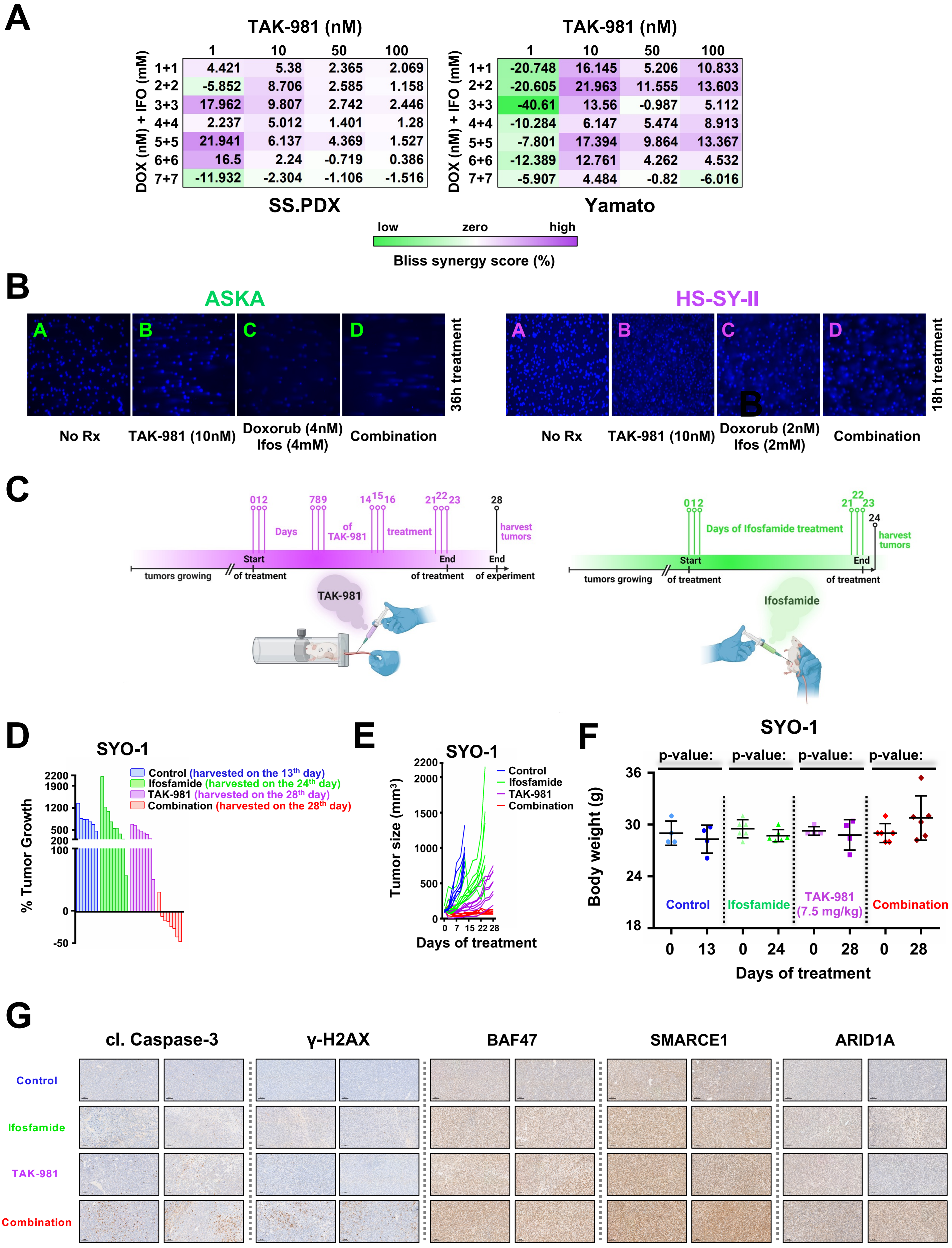
A
TAK-981 (nM)
1 10 50 100
DOX (nM) + IFO (mM)
1+1 4.421 5.38 2.365 2.069
2+2 -5.852 8.706 2.585 1.158
3+3 17.962 9.807 2.742 2.446
4+4 2.237 5.012 1.401 1.28
5+5 21.941 6.137 4.369 1.527
6+6 16.5 2.24 -0.719 0.386
7+7 -11.932 -2.304 -1.106 -1.516
SS.PDX
TAK-981 (nM)
1 10 50 100
DOX (nM) + IFO (mM)
1+1 -20.748 16.145 5.206 10.833
2+2 -20.605 21.963 11.555 13.603
3+3 -40.61 13.56 -0.987 5.112
4+4 -10.284 6.147 5.474 8.913
5+5 -7.801 17.394 9.864 13.367
6+6 -12.389 12.761 4.262 4.532
7+7 -5.907 4.484 -0.82 -6.016
Yamato
low zero high
Bliss synergy score (%)
B
ASKA
A B C D
36h treatment
No Rx
TAK-981 (10nM)
Doxorub (4nM) Ifos (4mM)
Combination
HS-SY-II
A B C D
18h treatment
No Rx
TAK-981 (10nM)
Doxorub (2nM) Ifos (2mM)
Combination
C
0 1 2 7 8 9 14 15 16 21 22 23 28
Days of TAK-981 treatment
harvest tumors
tumors growing
Start of treatment
End of treatment
End of experiment
TAK-981
0 1 2 21 22 23 24
Days of Ifosfamide treatment
harvest tumors
tumors growing
Start of treatment
End of treatment
Ifosfamide
D
SYO-1
% Tumor Growth
Control (harvested on the 13th day)
Ifosfamide (harvested on the 24th day)
TAK-981 (harvested on the 28th day)
Combination (harvested on the 28th day)
E
SYO-1
Tumor size (mm³)
Control
Ifosfamide
TAK-981
Combination
Days of treatment
F
SYO-1
p-value: p-value: p-value: p-value:
Body weight (g)
Control
Ifosfamide
TAK-981 (7.5 mg/kg)
Combination
Days of treatment
G
cl. Caspase-3
γ-H2AX
BAF47
SMARCE1
ARID1A
Control
Ifosfamide
TAK-981
Combination

### Figure S9

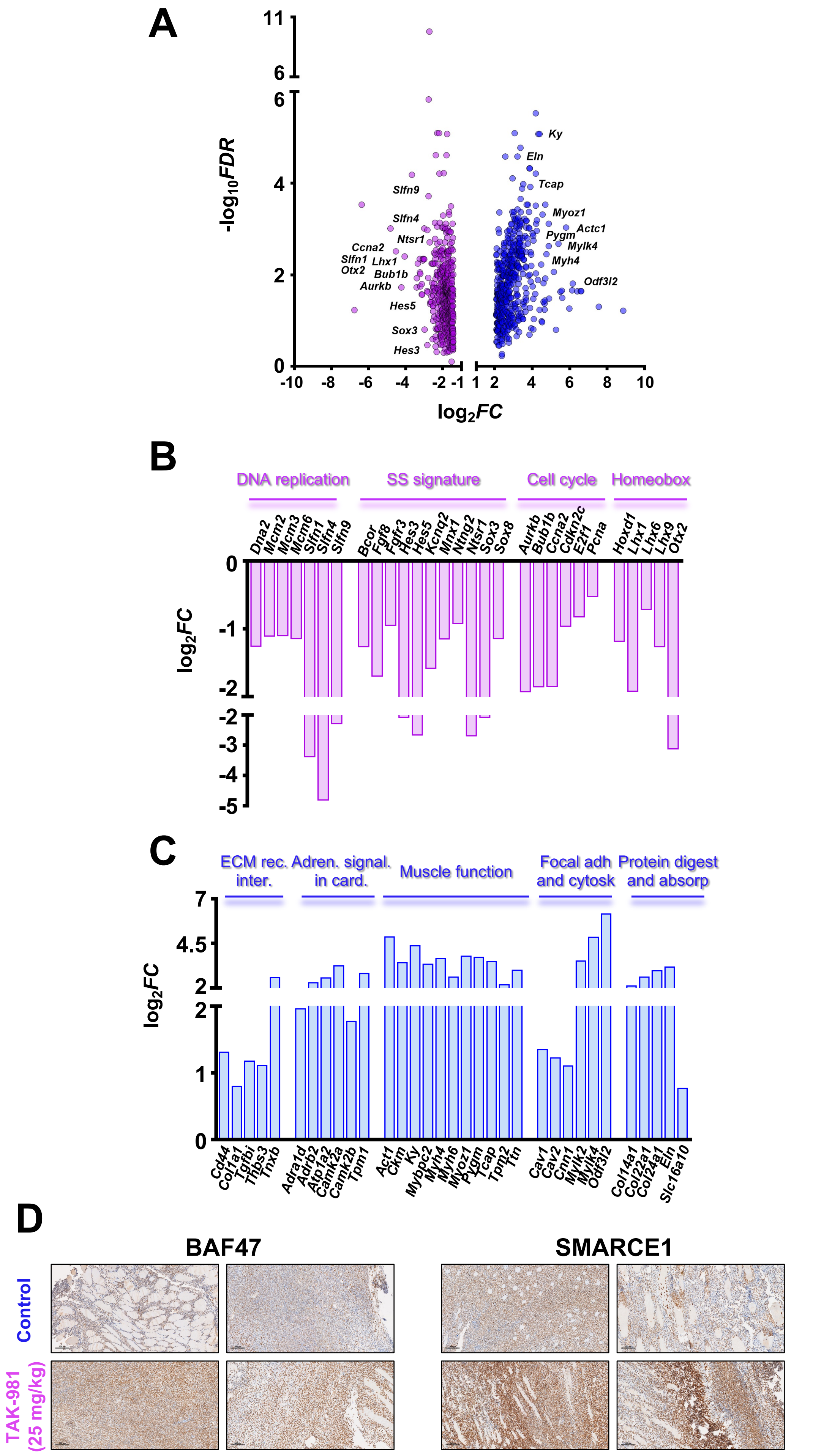
A
-log10FDR
log2FC
Ky
Eln
Tcap
Myoz1
Pygm
Actc1
Mylk4
Myh4
Odf3l2
Slfn9
Slfn4
Ntsr1
Ccna2
Slfn1
Lhx1
Otx2
Bub1b
Aurkb
Hes5
Sox3
Hes3
B
DNA replication
SS signature
Cell cycle
Homeobox
C
ECM rec. inter.
Adren. signal. in card.
Muscle function
Focal adh and cytosk
Protein digest and absorp
D
BAF47
SMARCE1
Control
TAK-981 (25 mg/kg)

Figure S10
